# Network-based prioritisation and validation of novel regulators of vascular smooth muscle cell proliferation in disease

**DOI:** 10.1101/2023.08.25.554834

**Authors:** Jordi Lambert, Sebnem Oc, Matthew D Worssam, Daniel Häußler, Nichola L Figg, Ruby Baxter, Kirsty Foote, Alison Finigan, Krishnaa T Mahbubani, Martin R Bennett, Achim Krüger, Mikhail Spivakov, Helle F Jørgensen

## Abstract

Aberrant vascular smooth muscle cell (VSMC) homeostasis and proliferation are hallmarks of vascular diseases causing heart attack and stroke. To elucidate molecular determinants governing VSMC proliferation, we reconstructed gene regulatory networks from single cell transcriptomics and epigenetic profiling. We find progressive activation of enhancers at disease-relevant loci in VSMCs that don’t respond to injury and proliferation-predisposed cells. Our analysis suggests that while many transcription factors are shared, their target loci differ between VSMC states. Comparison of rewiring between VSMC subsets and *in silico* perturbation analysis prioritised novel regulators, including RUNX1 and the network target TIMP1. We experimentally validated that the pioneer factor RUNX1 increased VSMC responsiveness and show that TIMP1 feeds back to promote VSMC proliferation through CD74-mediated STAT3 signalling. Both RUNX1 and the TIMP1-CD74 axis were expressed in hVSMCs, at low frequency in normal arteries and increased in disease, suggesting clinical significance and potential as novel vascular disease targets.

## Introduction

Vascular smooth muscle cell (VSMC) proliferation underlies cell accumulation in atherosclerotic vascular disease and restenosis that occurs in response to stenting and vascular grafting. While evidence from genome-wide association studies points to VSMCs as important hereditary determinants of cardiovascular disease (Örd *et al*, 2021; Turner *et al*, 2022; Liu *et al*, 2018), clinical targeting of VSMCs for these conditions is presently unexplored. VSMC lesion investment in experimental atherosclerosis results from extensive clonal expansion of a small number of cells (Chappell *et al*, 2016; Feil *et al*, 2014). Similar VSMC oligoclonality has been demonstrated in other vascular disease models using genetic lineage tracing of VSMCs (Clément *et al*, 2019; Chen *et al*, 2019), and clonal VSMC contribution in human disease has been proposed (Steffensen *et al*, 2022; Lin *et al*, 2023). Mapping of clone dynamics revealed that the oligoclonal contribution in atherosclerosis and following vascular injury results from activation of VSMC proliferation at low frequency, which is recapitulated in vitro (Worssam *et al*, 2022). The frequency of proliferating VSMC clones is affected by altered cell-cell communication (Misra *et al*, 2018) and is increased in ageing (Sheikh *et al*, 2018). This demonstrates that VSMC clonal proliferation is modifiable and suggests that changes in activation frequency could underlie increased vascular disease risk.

Single cell RNA-sequencing (scRNA-seq) studies in human atherosclerotic lesions and mouse disease models have shown remarkable transcriptional heterogeneity of VSMC-derived cells in disease (reviewed in (Winther *et al*, 2022)). VSMC can adopt states that range from quiescence, with high expression of contractile genes (such as *MYH11* and *ACTA2*), in healthy arteries, to cells that have induced signatures of other cell types, including fibromyocytes (e.g. *TNFRSF11B*), macrophages (*LGALS3*), and chondrocytes (*RUNX2*) in addition to extracellular matrix (ECM) proteins and remodellers that are characteristic of the so-called “synthetic” state generated by classical VSMC phenotypic switching (Pan *et al*, 2020; Wirka *et al*, 2019; Conklin *et al*, 2021; Dobnikar *et al*, 2018).

Functional genomics in lineage-traced animals have shed light on genetic determinants and mechanisms regulating VSMC cell state changes in disease (Örd *et al*, 2021; Turner *et al*, 2022; Wang *et al*, 2021). These novel experimental approaches have also revealed unexpected regulators of VSMC investment, such as the pluripotency factor OCT4 (Cherepanova *et al*, 2016), and identified a mesenchymal VSMC-derived state marked by expression of VCAM1 and SCA1 (Dobnikar *et al*, 2018; Pan *et al*, 2020). SCA1- positive VSMCs are found at low frequency in healthy arteries, increased in disease models and have been linked to VSMC priming and proliferation (Wang *et al*, 2020; Worssam *et al*, 2022; Dobnikar *et al*, 2018; Wirka *et al*, 2019). The existence of molecular heterogeneity of VSMCs prior to disease development may explain apparently distinct effects of VSMC regulators in different contexts (Yoshida *et al*, 2008; Owsiany *et al*, 2022). However, the events governing activation of clonal VSMC proliferation are understudied (Worssam & Jørgensen, 2021).

Recently, integration of single cell resolution gene expression with epigenetic information has demonstrated great potential for identification of factors regulating developmental processes and disease (Kamimoto *et al*, 2023). Here we use this approach to model regulatory networks upon acute vascular injury when VSMC proliferation initiates. We find differential use of transcription factors in distinct VSMC transcriptional states along a proliferation-associated trajectory, possibly explaining context-specific effects of VSMC regulators. The analysis identifies known and novel candidate regulators that are prioritised using *in silico* simulation analysis. We functionally implicate RUNX1 and TIMP1 in driving activation of VSMC proliferation and suggest these mechanisms also operate at early stages of human disease development.

## Results

### VSMC activation is associated with *de novo* chromatin opening at distal sites linked to vascular disease-associated genes

To investigate the molecular regulation of VSMC proliferation, we elicited an acute injury response by ligation of the left carotid artery. This model leads to reproducible VSMC phenotypic switching and initiation of VSMC proliferation 5-7 days after injury (Kumar & Lindner, 1997). Carotid ligation surgery was performed in Myh11-CreERt2, Rosa26-EYFP (Myh11-EYFP) animals after tamoxifen-induction of heritable EYFP expression in VSMCs to overcome the rapid loss of VSMC marker expression after injury. VSMCs expressing SCA1 have increased proliferative capacity (Dobnikar *et al*, 2018; Worssam *et al*, 2022; Pan *et al*, 2020). We therefore mapped chromatin accessibility changes using bulk ATAC-seq for SCA1+ and SCA1- lineage traced, EYFP+ VSMCs from injured animals separately, and compared these to EYFP+ VSMCs from littermate control animals that had not undergone surgery. Samples from all three conditions (SCA1+, SCA1- and Control) had high signal-to-noise ratio and strong correlation of peak intensity across replicates (**Fig. 1A, Extended Data Fig. S1A-C**).

**Figure 1.**
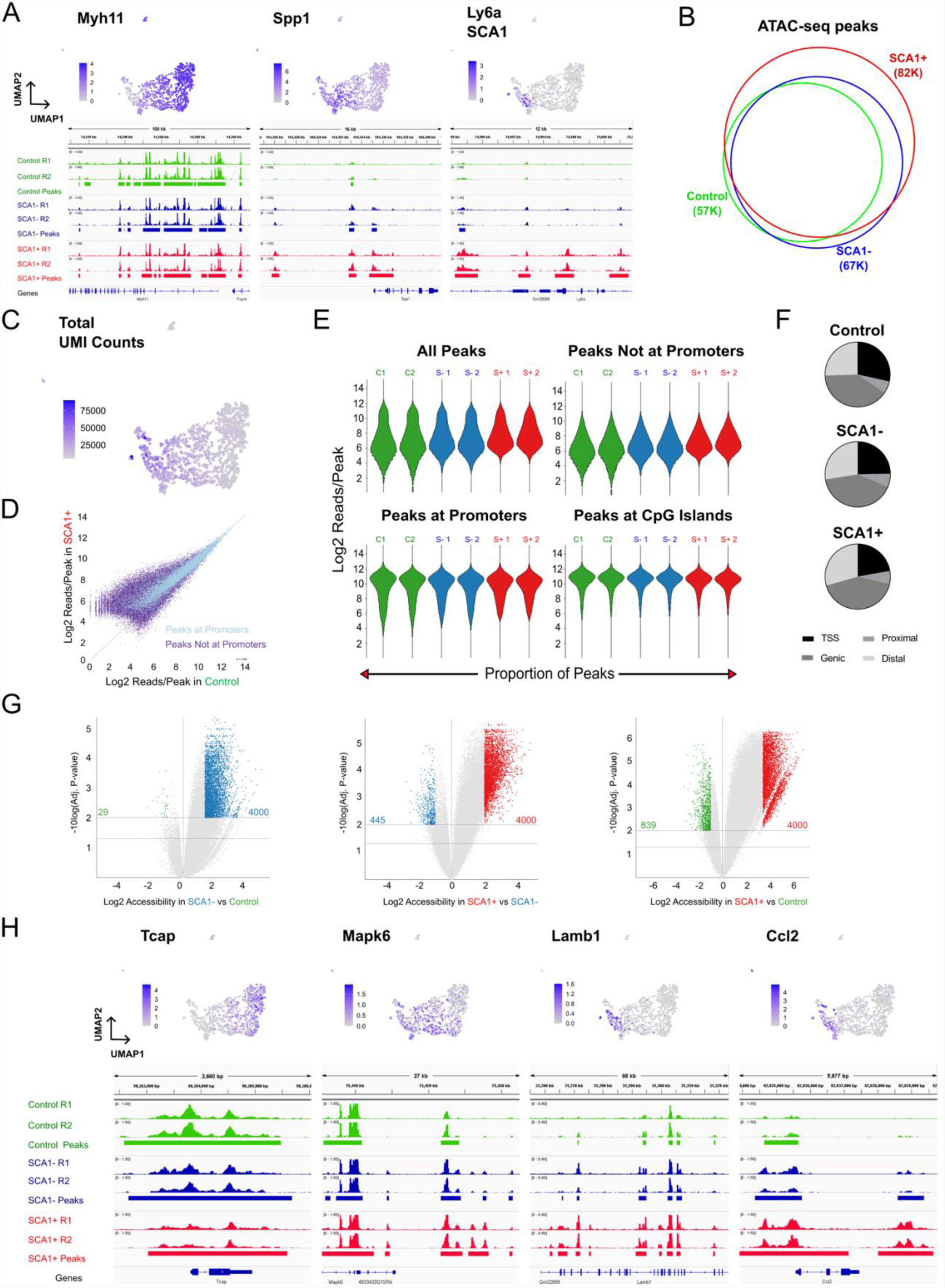
Activation of VSMCs results in widespread epigenetic activation of distal elements relevant for VSMC function and disease. **(A)** ATAC-seq traces for markers of contractile VSMCs (*Myh11*), the synthetic phenotype (*Spp1*) and a VSMC transition state (*Ly6a*, encoding SCA1) in EYFP+ lineage-traced VSMCs isolated from healthy control Myh11-EYFP animals (green), and either EYFP+SCA1-(SCA1-, blue) or EYFP+SCA1+ (SCA1+, red) cells isolated from injured arteries. Aligned reads for two independent experiments and reproducible peaks (horizontal bars) are shown. UMAPs of scRNA-seq profiles of VSMC-derived cells 7 days after injury (<u>GSE162167</u>) showing associated gene expression levels across (top). (**B**) Venn diagram representing overlap of ATAC-seq peaks in the three sample types. (**C**) UMAP of VSMC scRNA-seq data described in A, showing (<u>GSE162167</u>) the number of detected transcripts (unique molecular identifiers, UMI) per cell. (**D**) Scatterplot showing read density in healthy VSMCs (x-axis) and SCA1+ VSMCs after injury (y-axis) for ATAC-seq peaks at promoters (light blue) and peaks that are not at promoters (purple). (**E**) Violin plots showing distribution of peak read densities in each sample for all peaks, or only non-promoter, promoter and CpG island peaks. (**F**) Pie charts showing the proportion of peaks within 1 kb of transcriptional start sites (TSS), at promoter proximal regions (proximal), within genes (genic) or at intergenic elements (distal). (**G**) Volcano plots showing fold change of peak intensity between indicated samples. Horizontal lines indicate significance thresholds of q<0.01 (top) and q<0.05 (bottom). Differentially accessible peaks used for pathway enrichment analysis are highlighted (fold change>1.5 and of q<0.01, or top 4000 ranked by fold change). (**H**) Gene expression and ATAC-seq data tracks for genes showing higher accessibility in control samples (*Tcap*) or SCA1+ cells after injury (*Mapk6*, *Lamb1*, *Ccl2*).

No major changes in accessibility were detected at *Myh11* and other contractile genes that are downregulated after injury (**Fig. 1A**), consistent with the documented retention of the active H3K4me2 mark at contractile genes in the synthetic state (McDonald *et al*, 2005; Gomez *et al*, 2013). Genes that are induced in most VSMCs after injury (e.g., *Spp1*) displayed increased accessibility in both SCA1+ and SCA1- VSMCs, whereas the chromatin at the SCA1-encoding *Ly6a* locus became more accessible selectively in SCA1+ cells (**Fig. 1A)**. A total of 84K peaks were detected across the three conditions and 96% (82K) of these were accessible in SCA1+ cells (**Fig. 1B**). This progressive genome-wide *de novo* chromatin opening in SCA1+ (82K peaks), SCA1- (67K) compared to control samples (57K), reflected the increased transcription observed by scRNA-seq in *Ly6a*+ cells after injury (**Fig. 1C**).

The intensity of ATAC-seq signals was also generally increased in SCA1+ samples compared to control VSMCs (**Fig. 1D**). Increased peak intensity was most obvious at regions with lower accessibility, which is a characteristic of enhancers. Consistently, ATAC-seq peak intensity was shifted upwards for non-promoter peaks, but not changed for promoters and CpG islands (**Fig. 1E**). The proportion of peaks at distal elements was also higher in SCA1+ cells, compared to both SCA1- and control samples (**Fig. 1F**), confirming enhancer activation. Differential accessibility analysis across all 84K peaks, confirmed significantly higher signal intensity at many peaks in SCA1+ samples, compared to both SCA1- (14,254 peaks) and control VSMCs (28,303 peaks, **Fig. 1G**). Much fewer peaks were more accessible in controls compared to VSMCs from injured arteries (839 compared to SCA1+, and 29 compared to SCA1-), and associated genes (e.g. *Tcap,* **Fig. 1H**) were enriched for muscle contraction gene ontology (GO) terms (**Extended Data Fig. S1D**). In contrast, genes associated with the regions with the most increased chromatin accessibility in SCA1+ compared to control samples were enriched for vascular diseases, including atherosclerosis, and biological process ontology terms linked to VSMC proliferation, such as regulation of signal transduction (*Mapk6*), inflammation (*Ccl2*) and cell adhesion (*Lamb1*)(**Fig. 1H, Extended Data Fig. S1E-G**). This analysis demonstrates that injury-induced VSMC activation results in opening of chromatin at enhancers of genes related to the functions of synthetic VSMCs and vascular disease, and that this is most pronounced in VSMCs expressing SCA1.

### Rewiring of shared factors between injury-induced VSMC states

In order to identify factors that control activation of VSMC proliferation, we wanted to compare how gene expression is orchestrated in VSMCs that respond to injury versus those that do not. We used scRNA-seq profiles of VSMCs analysed at the very onset of VSMC proliferation (5 days post injury), where we previously identified a proliferation-associated trajectory (Worssam *et al*, 2022). Based on their position along this trajectory, we classified VSMCs as “non-responding” (“*Non-RSP*”, defined by persistent high levels of contractile gene expression), a “linking” cell population (“*LNK*”), leading on to a pre-proliferative state including SCA1-expressing cells (“*PrP*”) and, finally, actively cycling cells that express *Mki67* and have high G2/S scores (“*CYC*”) (**Fig. 2A, Extended Data Fig. S2A** and (Worssam *et al*, 2022)).

**Figure 2.**
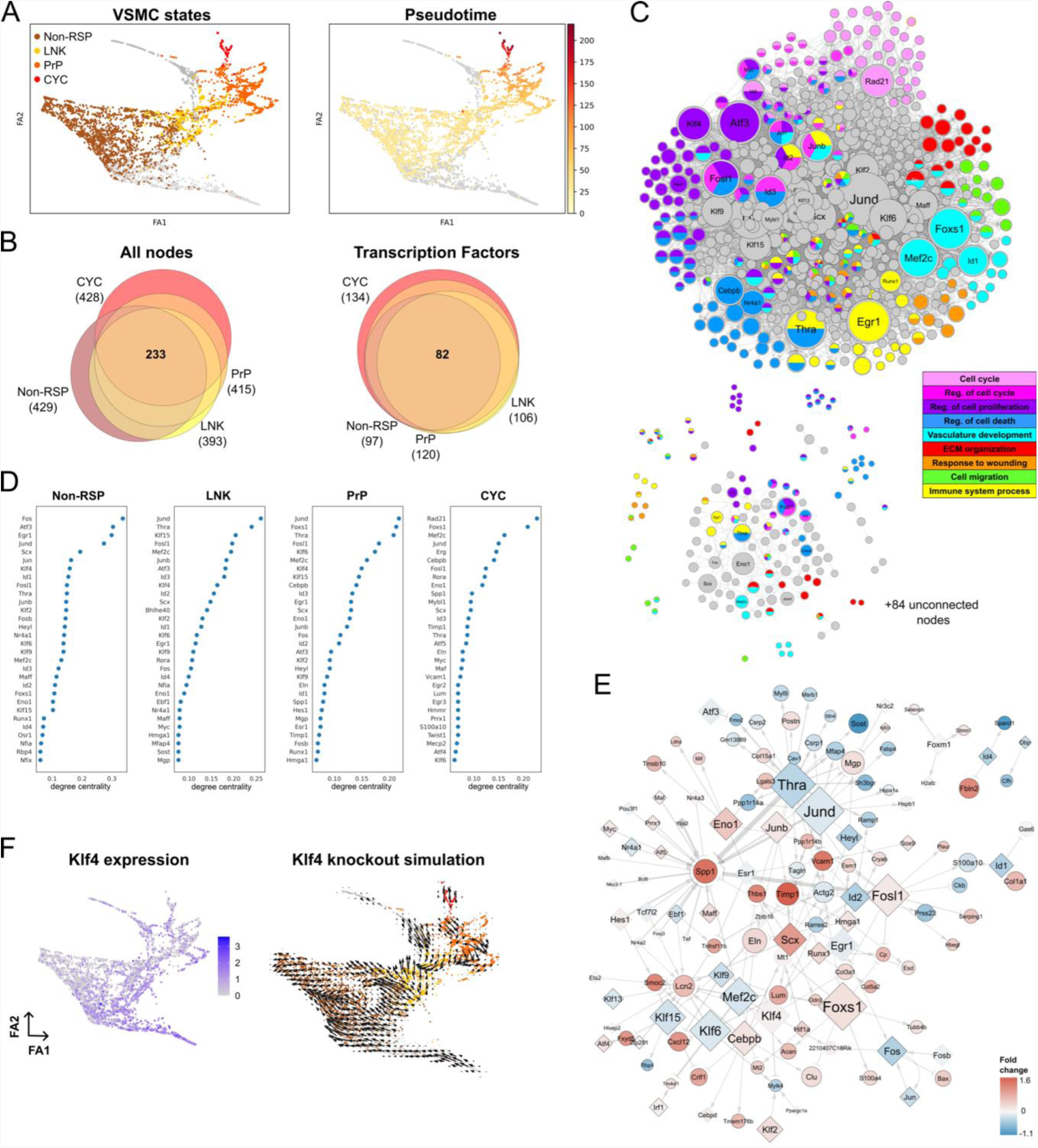
VSMC state-specific gene regulatory networks. **(A)** VSMC states (left; Non-RSP: brown, LNK: yellow, PrP: orange and CYC: red, non-assigned: grey) or proliferation-associated pseudotime (right, white-to-red colour scale) are indicated on force-directed graph representation of scRNA-seq data from VSMC-derived cells isolated 5 days after injury (<u>GSE162167</u>). **(B)** Euler diagrams for all nodes (left) and transcription factors (right) in the four gene regulatory networks (GRNs) colour-coded as in panel A. The total number of nodes or transcription factors are shown in brackets. **(C)** The union (top) and intersection (lower panel) of Non-RSP, LNK, PrP, and CYC networks coloured by associated GO terms. The node size reflects degree centrality. **(D)** Nodes with top (30) degree centrality scores in each GRN. **(E)** The PrP network including interactions with connectivity threshold above 0.1. Differential expression between PrP and Non-RSP cells is indicated by a blue (higher in Non-RSP) to red (higher in PrP) colour scale. Black borders indicate significantly differential expression (p_adj_ <0.05). Edge widths reflect edge connectivity. **(F)** Force-directed graph projections showing gene expression (left) and *in silico* knockout simulation (right) for Klf4.

To construct gene regulatory networks (GRNs) for these four VSMC states, we used CellOracle (Kamimoto *et al*, 2023); we first generated a VSMC-specific transcription factor-gene interaction network based on all 84K ATAC-seq peaks, which was combined with gene expression profiles for cells in each of the VSMC states described above. The resulting four GRNs for Non-RSP, LNK, PrP and CYC VSMCs had comparable centrality score distribution (**Extended Data Fig. S2B**). We found a substantial node overlap between the networks, with 233/597 (37%) of all identified genes, including 82/140 (59%) transcription factors, shared between the four GRNs (**Fig. 2B, Supplementary Table S1AB**). Despite the high number of common nodes, few interactions were present in all four GRNs (**Fig. 2C, lower panel**). Accordingly, abundant changes in the network topology rankings, such as degree centrality of transcription factors, were observed between networks (**Fig. 2D, Supplementary Table S1C**). GRN modelling using the scRNA-seq profiles of VSMCs analysed 7 days after injury was conducted in parallel and produced comparable results (**Extended Data Fig. 2D**).

Overall, the network nodes were enriched for processes related to VSMC biology or vascular injury, including ECM organisation, actin organisation, response to stimuli and regulation of cell proliferation, (**Fig. 2C, Supplementary Table S1B**). Generally, specific regulatory circuits driving biological processes were not detected (**Extended Data Fig. S2C)**. However, the CYC network included a substructure consisting of cell cycle-associated genes regulated by RAD21, a subunit of the cohesion complex that has roles in chromosome segregation, genome stability and transcriptional regulation (Cheng *et al*, 2020) (**Extended Data Fig. S2B, C**). The CYC network, which had the most unique factors (14 transcription factors, **Fig. 2B**), was dominated by genes driving cell cycle progression, such as Myc, Mybl1, Erg and Rad21 (**Fig. 2D**). To study processes and factors driving activation, rather than progression, of VSMC proliferation we therefore focussed on comparison of the PrP network, shown in **Fig. 2E**, to that of Non-RSP cells.

The generated GRNs enabled computational simulation of how perturbation of transcription factor levels would affect VSMC state. KLF4 is a known regulator of VSMC phenotypic switching that promotes a macrophage-like state in atherosclerosis (Shankman *et al*, 2015) and the *Klf4* node had high centrality in all networks apart from that of proliferating cells (CYC). *In silico* simulation of KLF4 depletion predicted a general shift towards the non-responding cell state, concomitantly with promotion of CYC cells (**Fig. 2F**). This context specific effect of KLF4 depletion is consistent with the phenotype of VSMC-specific KLF4 knockout animals after injury, which show delayed downregulation of contractile genes, but accelerated neointima formation (Yoshida *et al*, 2008). *In silico* simulations of other factors with high connectivity scores in all networks also suggested an impact of central transcription factors on VSMC state. For example, MEF2C had high degree centrality score in all four networks, but higher “betweenness” score in the Non-RSP and LNK GRNs (**Fig. 2D, Supplementary Table S1C**). Consequently, *in silico* simulations predicted movement away from the Non-RSP state that maintains contractile gene expression (**Extended Data Fig. S3),** which aligns with a reported role of MEF2C in driving VSMC differentiation (Pagiatakis *et al*, 2012). Thyroid hormone receptor alpha (THRA) that has been suggested to impact VSMC cholesterol metabolism (Neggazi *et al*, 2018), was also a central node in all networks (**Fig. 2D**). *Thra* is expressed at lower levels in PrP compared to Non-RSP VSMCs and negatively interacts with synthetic genes such as *Spp1*, *Vcam1* and *Mgp* in the PrP network, while promoting cytoskeletal genes (*Actg2*, *Pdlim4*, *Cdc42ep3*) in the Non-RSP GRN (**Supplementary Table S1A**). This suggests that THRA may act to safeguard the contractile state, as also indicated by simulation results (**Extended Data Fig. S3**), thereby protecting from disease-associated phenotypic switching. Overall, this analysis suggests that context-specific differences in transcription factor-target gene interactions contributes to driving activation of VSMC proliferation and highlights both known and putative novel factors controlling VSMC state.

### Comparison of VSMC state-specific GRNs identifies candidate regulators of VSMC activation

The GRN analysis presented above suggested that differential use of shared factors plays a central role in VSMC activation. To assess whether this regulatory rewiring between VSMC states is relevant for regulating gene expression in humans, we used published scRNA-seq profiles of human carotid plaque cells (Pan *et al*, 2020). We selected mural cells from this dataset based on expression of contractile and synthetic VSMC markers and clustered these into 6 groups (**Fig. 3A, Supplementary Table S2**). Clusters 0-2 had abundant contractile gene expression, cluster 3 cells expressed *TNFRSF11B* that was identified as a fibromyocyte marker (Wirka *et al*, 2019), *VCAM1* was expressed in clusters 3 and 4, and cluster 5 had increased levels of ribosomal proteins (**Extended Data Fig. S4**). We organised cells by these clusters to visualise how the state of human plaque VSMCs expression correlate with the effects of transcriptional rewiring after injury. Genes showing highest rewiring between the Non-RSP and PrP GRNs and their direct interactors were hierarchically clustered by expression in post-injury murine VSMCs and side-by side comparison revealed a similar expression pattern in the human plaque VSMCs (**Fig. 3B**), indicating that the GRN analysis identify genes that are implicated in atherosclerosis in human arteries.

**Figure 3.**
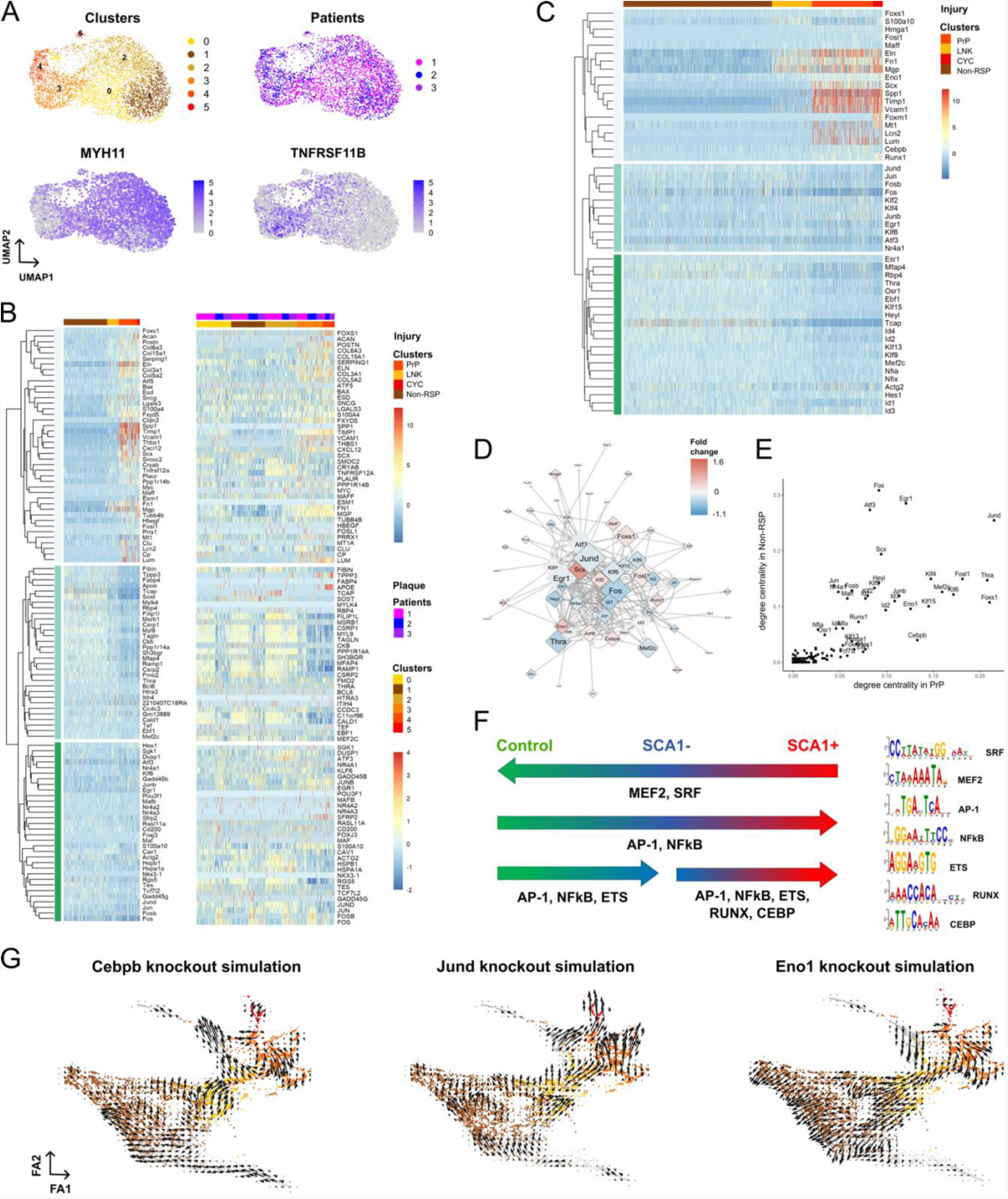
Rewiring of GRNs across cell states identifies candidate regulators of VSMC activation. **(A)** UMAPs showing cell clusters, patients, and expression of selected genes in VSMCs from a human carotid plaque dataset (Pan *et al*, 2020). **(B)** Heatmap showing expression of genes ranked in the top 10 according to rewiring score between PrP and Non-RSP GRNs and their direct interactors in VSMCs analysed 5 days after injury (left) or in human carotid plaque VSMCs (Pan *et al*, 2020). Genes are clustered based on expression along the mouse VSMC states. **(C)** Heatmap showing expression of the genes with the 50 highest rewiring scores between PrP and Non-RSP GRNs in the mouse dataset. **(D)** Transcription factor subnetwork for the union of PrP and Non-RSP networks, coloured by differential expression as in Figure 2E. **(E)** Comparison of out-degree centrality scores in PrP and Non-RSP GRNs for the top scoring genes in either network. **(F)** Summary of motif enrichment analysis for peaks showing higher accessibility relative to all peaks for the indicated comparisons (left). Detected motifs are shown on the right. **(G)** Force-directed graphs showing the result of knockout simulation results for the selected genes.

Genes showing most change between the Non-RSP and PrP networks were predominantly transcription factors (36/50) and fell into three groups with different expression patterns (**Fig. 3C**). Group 1 increased expression along the injury-associated trajectory, including VSMC phenotypic switching marker genes (*Spp1*, *Fn1*, *Mgp*), *Timp1* and *Vcam1*, that marks activated VSMCs together with SCA1/*Ly6a* (Pan *et al*, 2020). This group also comprised transcription factors *Scx*, *Cebpb*, *Runx1* and *Eno1*. Group 2 had clearly reduced expression, including *Mef2c, Thra, Hes*, *Heyl,* Kruppel like factors (KLF) 9, 13, and 15, and several inhibitor of differentiation (Id) genes. The third group of genes showed little variation in expression between the non-responding and proliferation-associated states. This group contains factors reported to affect VSMC regulation, e.g. KLF4 (Shankman *et al*, 2015; Yoshida *et al*, 2008), EGR1 (Wada *et al*, 2003), ATF3 (Wang *et al*, 2021), and AP-1 subunits (Zhao *et al*, 2019). Transcription factors generally showed less differential expression between PrP and Non-RSP VSMCs compared to target genes (**Fig. 3D** *vs.* **Fig 2E**). In contrast, substantial topology score differences were observed (**Fig. 3E**), illustrating the added insight provided by the GRN approach.

We also assessed whether the increased chromatin accessibility in activated VSMCs is related to specific transcription factors. Motif enrichment analysis for ATAC-seq peak regions that had differential accessibility between conditions is summarised in **Fig. 3F**. The MEF2 motif was enriched along with SRF for peaks showing higher accessibility in control samples compared to SCA1+ VSMCs from injured arteries. Peaks that gained *de novo* accessibility after injury were enriched for AP-1, NFκB, CEBP, ETS and RUNX binding sites. Interestingly, RUNX, NFκB and CEBP motifs were also detected together with AP-1 binding sites (**Extended Data Fig. S5A**), suggesting co-regulatory interaction at some loci.

We then performed *in silico* deletion experiments for the candidate VSMC regulators identified by the GRN analysis, similarly to the analysis presented above. Like for KLF4, we found that several factors had context-specific functions (**Fig. 3G, Extended Data Fig. S3**). This includes the ETS target Egr1 and Cebpb/d, which have previously been implicated in VSMC regulation (Ackers-Johnson *et al*, 2015). The impact of AP-1 on VSMC activation was predicted to be subunit-dependent; while simulation suggested that *Fos*, *Jun* or *Jund* knockout would promote VSMC activation, loss of *Junb* was predicted to inhibit transition to the PrP state (**Fig. 3G, Extended Data Fig. S3**). In contrast, *in silico* depletion of the runt-related transcription factor *Runx1* or *Eno1*, which encodes both the glycolytic enzyme alpha-enolase and a nuclear protein previously named Myc-binding protein, was predicted to block progression towards proliferation (**Extended Data Fig. S3**). Taken together this analysis identified novel transcription factors predicted to control activation of VSMC proliferation and highlights the benefit of GRN studies in prioritisation of regulators relative to analyses based solely on differential expression.

### RUNX1 promotes VSMC proliferation

The analyses above highlighted the RUNX motif, which was also identified in relation to VSMC regulation by single cells studies in experimental atherosclerosis (Örd *et al*, 2021). The family member RUNX2 is known to induce expression of ECM genes, which stimulates VSMC calcifications in lesions, and has been proposed to regulate cell proliferation (Komori, 2019; Durham *et al*, 2018). However, *Runx2*, as well as *Runx3*, was lowly expressed at this stage of VSMC activation (**Extended Data Fig. S2A**). In contrast, *Runx1* transcripts were detected at higher levels in the PrP and CYC states compared to non-responding cells (**Fig. 4A**) and increased chromatin accessibility was detected at both *Runx1* promoters in SCA1+ VSMCs compared to the control samples (**Extended Data Fig. S5B**). We therefore further investigated how RUNX1, which is a key regulator in haematopoiesis, affects VSMC activation.

**Figure 4.**
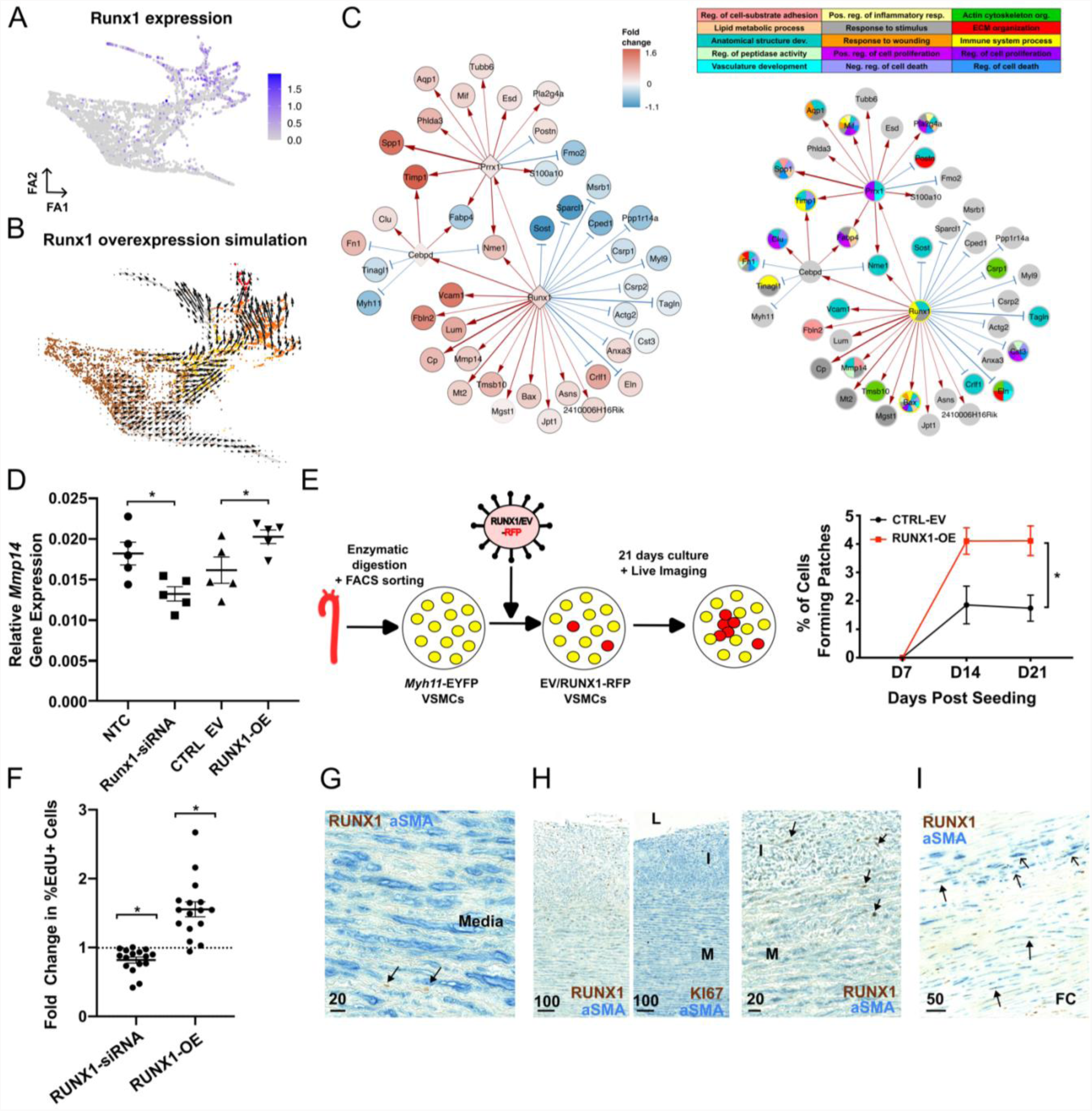
RUNX1-mediated regulation of VSMC proliferation. **(A, B)** Force-directed graph of VSMCs isolated 5 days after injury showing *Runx1* expression **(A)** or the result of RUNX1 overexpression simulation **(B)**. **(C)** Direct and indirect targets of Runx1 in the PrP GRN showing differential expression between PrP and Non-RSP cells on a blue (higher in Non-RSP) to red (higher in PrP) colour scale. Black borders indicate significantly differential expression (p_adj_<0.05). Edge widths reflect edge connectivity and edge colour show positive (red) and negative (blue) interactions. **(D)** *Mmp14* transcript levels detected by quantitative RT-PCR in mVSMCs transfected with non-targeting (NTC) or *Runx1*-targeting siRNA (Runx1-siRNA), or transduced with an empty vector (CTRL-EV) or RUNX1-overexpressing (OE) lentivirus. Dots represent cells from independent animals (N=5) and lines indicate mean expression. **(E)** Schematic of clonal proliferation assay where a red fluorescent protein (RFP)-expressing lentivirus is used to test the effect of RUNX1 cDNA relative to an empty control vector (EV) on the ability of lineage-labelled VSMCs from Myh11-EYFP animals to form colonies (left). Right panel shows the percentage of cells forming a clonal VSMC patch in RUNX1-OE and empty vector control cells (CTRL-EV). **(F)** Fold change in %EdU+ cells after siRNA-mediated RUNX1 depletion (RUNX1-siRNA, relative to non-targeting siRNA-treated cells) and lentivirus-mediated RUNX1-overexpression of hVSMCs (RUNX1-OE, relative to cells transduced with empty vector virus). **(G-I)** Representative immunostaining of non-plaque aorta (G and H, N=10) or carotid endarterectomy samples (I, N=6) for RUNX1, αSMA and KI67 as indicated. Scale bars: G: 20 µm, H: 100 µm (left, middle) and 20 µm (right), I: 50 µm. FC: fibrous cap, L: Lumen, I: Intima, M: Media. Examples of RUNX1+ (closed arrowheads) and RUNX1-cells (open arrowheads) that are αSMA+ are indicated. Asterisk indicates p<0.05.

GRN-based simulation of elevated RUNX1 expression predicted that RUNX1 promotes the transition towards VSMC proliferation (**Fig. 4B**), as expected from the limitation of proliferation observed when simulating *Runx1*-knockout (**Extended Data Fig. S3**). Consistent with this idea, the PrP GRN predicts that RUNX1 upregulates *Vcam1*, *Lum* and *Mmp14* and other injury-induced genes. Gene ontology analysis of all direct and indirect RUNX1 target genes suggests that RUNX1 could affect cell-substrate adhesion (*Fbln2, Mmp14*), response to stimulus (*Cp*, *Mt2*) and ECM organisation (*Eln*) (**Fig. 4C**). Interestingly, four of the 15 direct RUNX1 target genes were previously associated with activated, SCA1+ VSMCs in healthy arteries (Dobnikar *et al*, 2018). We also found that expression of the predicted RUNX1 target, MMP14 is a hallmark of SCA1+ VSMCs (**Extended Data Fig. S5C**), further indicating that RUNX1 supports a state that is primed for proliferation.

We experimentally validated RUNX1-mediated regulation of predicted direct RUNX1 target genes *Mmp14* and *Cebpd*. Lineage-traced VSMCs were isolated from mouse aortas and cultured for three days to achieve culture-induced VSMC activation, before siRNA-mediated depletion of *Runx1*, which significantly reduced *Mmp14* expression (**Fig. 4D, Extended Data Fig. S5D**). Conversely, levels of both *Cepbd* and *Mmp14* were increased after lentiviral-delivery of RUNX1-encoding cDNA, compared to control cells (**Fig. 4D, Extended Data Fig. S5D**). To directly test whether RUNX1 affects activation of VSMC proliferation, we adapted an *in vitro* proliferation assay that reproduces the low frequency clonal VSMC expansion observed in vascular disease models (Worssam *et al*, 2022). Infection of lineage-labelled VSMCs with RFP-expressing lentivirus at low titre was followed by live cell imaging over three weeks to identify expanding clonal VSMC patches. Quantification of RFP/EYFP double positive clones showed a significantly increased clonal patch-forming ability for cells receiving lentivirus encoding RUNX1 compared to a control empty vector (**Fig 4E**).

In human VSMCs (hVSMCs), RUNX1-depletion decreased the percentage of EdU+ cells and, *vice versa*, overexpression of RUNX1 increased the percentage of EdU+ cells, relative to their respective controls. This suggests that stimulation of VSMC proliferation by RUNX1 is conserved in human (**Fig 4F**). To further assess RUNX1 functions in human disease, we immunostained arteries from organ donors that did not have detectable plaques but displayed varying degrees of intimal thickening, as well as atherosclerotic plaques from carotid endarterectomy. RUNX1 protein was detected infrequently in alpha smooth muscle actin (αSMA)+ cells in arteries with a healthy, organised morphology of the medial layer, but a higher proportion of medial RUNX1+αSMA+ cells were observed in arteries with evidence of perturbation (**Fig 4G**). In arteries displaying intimal thickening and proliferation (KI67), many αSMA+ cells within the intima expressed RUNX1 (**Fig 4H, Extended Data Fig. S5E**). In atherosclerotic lesions, αSMA+RUNX1+ cells were observed in the fibrous cap region but substantial heterogeneity was observed (**Fig 4I**). Jointly, these results demonstrate that RUNX1 regulates gene expression in VSMCs and promotes their proliferation. RUNX1 expression by VSMCs in human arteries suggests that it is implicated in dysregulation of proliferation in disease.

### TIMP1 promotes VSMC proliferation in an MMP-independent manner

Next, we used the GRN analysis to investigate how targets of the identified transcriptional rewiring could impact VSMC biology. To this end, we focussed on genes with high target gene score in the PrP or CYC regulatory networks (“degree centrality in”, **Supplemental Data 1**) that are expressed at higher levels in the PrP compared to the Non-RSP state. Pathway analysis highlighted gene ontology terms including ECM organization (*Spp1*, *Vcam1*, *Cxcl2*, *Postn*) and regulation of cell adhesion (*Eln*, *Lum*, *Fbln2*). We also found modulators of proteolysis (*Timp1*, *Serpine1*, *Thbs1*) among the targets of rewired transcription factors. This is consistent with a role of activated VSMCs in remodelling the cellular environment. We focussed on TIMP1 (Tissue Inhibitor of Metalloproteinases-1), which itself had high rewiring score, was an indirect target of RUNX1 and also received input from Cebpd, Myc, Prrx1 and Scx (**Fig. 3C, 4C, 5A, Extended Data Fig. S5E**). TIMP1 is both an inhibitor of matrix metalloproteinases (MMPs) and functions as a cytokine to, for example, induce fibroblast proliferation in cancer (Song *et al*, 2016; Schoeps *et al*, 2023). Furthermore, TIMP1 expression is upregulated by VSMCs in human and mouse atherosclerosis scRNA-seq datasets (**Extended Data Figs. S2A, S4** and (Dobnikar *et al*, 2018; Pan *et al*, 2020; Örd *et al*, 2021)). While increased TIMP1 protein levels have been associated with cardiovascular disease and severity (Silence *et al*, 2002; Sundström *et al*, 2004), the responsible cellular and molecular mechanisms are unknown.

**Figure 5.**
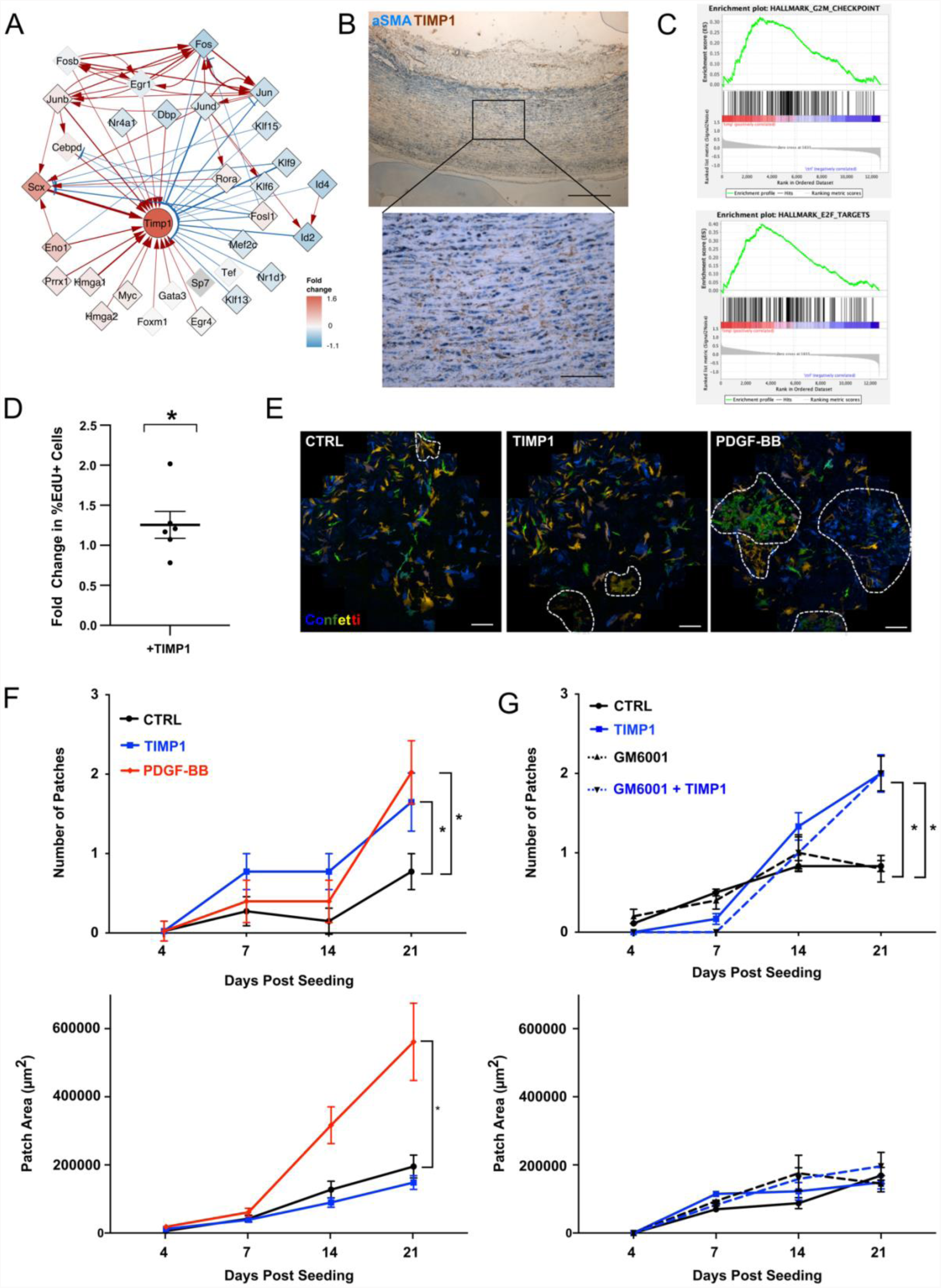
GRN analysis identifies TIMP1 as a functional gene target and driver of VSMC proliferation. **(A)** Predicted inputs to TIMP1 in the PrP GRN, coloured as described in left panel of Fig. 4C. **(B)** Representative immunohistochemistry image for αSMA (blue) and TIMP1 (brown) in non-plaque human aorta (N=7), scale bar=500 µm (overview) and 100 µm (zoomed view). **(C)** Enrichment plots from gene set enrichment analysis (GSEA) for E2F targets (p_adj_= 0.005) and G2M checkpoint genes (p_adj_= 0.04) in bulk RNA-seq data from human VSMCs (hVSMCs) treated with 500 ng/mL recombinant human (rh) TIMP1 for 6 hours versus control cells (N=6 independent hVSMC isolates). **(D)** Fold change in %EdU+ hVSMCs following 16 hours EdU incorporation in cells treated with 500 ng/mL rhTIMP1 relative to vehicle controls (N=6 independent hVSMC isolates). **(E)** Representative images of lineage labelled VSMCs isolated from Myh11-Confetti aortas and cultured for 21 days in the presence of vehicle control, 500 ng/mL recombinant murine TIMP1 or 2 ng/mL PDGF-BB. Scale bar=500 µm. **(F, G)** Quantification of number and size of clonally expanded patches of Confetti+ VSMCs over 21 days of culture. Statistical significance assessed via a generalised linear model, N=4 animals.

Immunostaining showed TIMP1 expression in αSMA+ cells located in the medial layer of non-plaque human arteries (**Fig. 5B, Extended Data Fig. S6A**), suggesting that it could affect VSMCs at an early disease stage in humans. To investigate the potential impact of TIMP1 protein on VSMC function, we treated six independent isolates of human VSMCs with recombinant TIMP1 protein and compared gene expression profiles to control samples using bulk RNA-seq. Gene set enrichment analysis showed significant induction of cell cycle genes (E2F targets and G2/M checkpoint) as well as fatty acid metabolism and oxidative phosphorylation gene sets (**Fig. 5C, Extended Data Fig. S6B**) in TIMP1-treated cells. Altered metabolism was suggested as an important determinant of VSMC phenotypic switching and is linked to proliferation (Newman *et al*, 2021). Consistent with an effect on cell cycle genes, a significant increase in %EdU+ cells was observed in TIMP1-treated hVSMCs versus controls (**Fig. 5D**). In the clonal proliferation assay, TIMP1-treatment also stimulated clonal expansion of VSMCs that were freshly isolated from lineage-labelled mouse aortas, similarly to PDGF (**Fig. 5E, F**). Both TIMP1 and PDGF increased VSMC clone formation frequency (**Fig. 5F**). However, in contrast to PDGF, TIMP1 did not change the size of clonal patches (**Fig. 5F, lower panel**), suggesting that the establishment and growth of VSMC clones are regulated independently. To test whether the ability of TIMP1 to promote VSMC proliferation is linked to inhibition of proteolysis, we added a broad-spectrum MMP inhibitor (GM6001). No effect of GM6001 on clone dynamics was observed in control nor in TIMP1-treated cells (**Fig. 5G**). Taken together, these results suggest that TIMP1 stimulates VSMC proliferation in a manner that is independent on its ability to inhibit MMP activity.

### TIMP1 induces STAT3 phosphorylation to induce VSMC proliferation

Multiple signalling pathways have been reported to be activated by TIMP1 in different contexts (Grünwald *et al*, 2019; Schoeps *et al*, 2023). To identify how TIMP1 promotes VSMC proliferation we therefore screened for phospho-kinase activation, which revealed increased levels of STAT3 phosphorylated at S727 in response to TIMP1 treatment of hVSMCs (**Fig. 6A, Extended Data Fig. S7A**). TIMP1-induced phospho-STAT3 (pSTAT3) was confirmed by western blotting in five independent hVSMC isolates, which showed increased phosphorylation at both S727 and T705, whereas total STAT3 protein levels were unchanged (**Fig. 6B, C**). The induction of STAT3 phosphorylation was observed within 5 minutes of TIMP1 addition and was transient, suggesting a direct response to TIMP1 (**Fig. 6B, C**). Phosphorylation of STAT3 leads to dimerization, nuclear import and binding at target gene promoters (Tesoriere *et al*, 2021). Consistent with its activation, STAT3 chromatin immunoprecipitation showed increased binding in TIMP1-treated cells, specifically at promoters of canonical STAT3 target genes (JUNB, TWIST1), compared to control cells (**Fig. 6D**).

**Figure 6.**
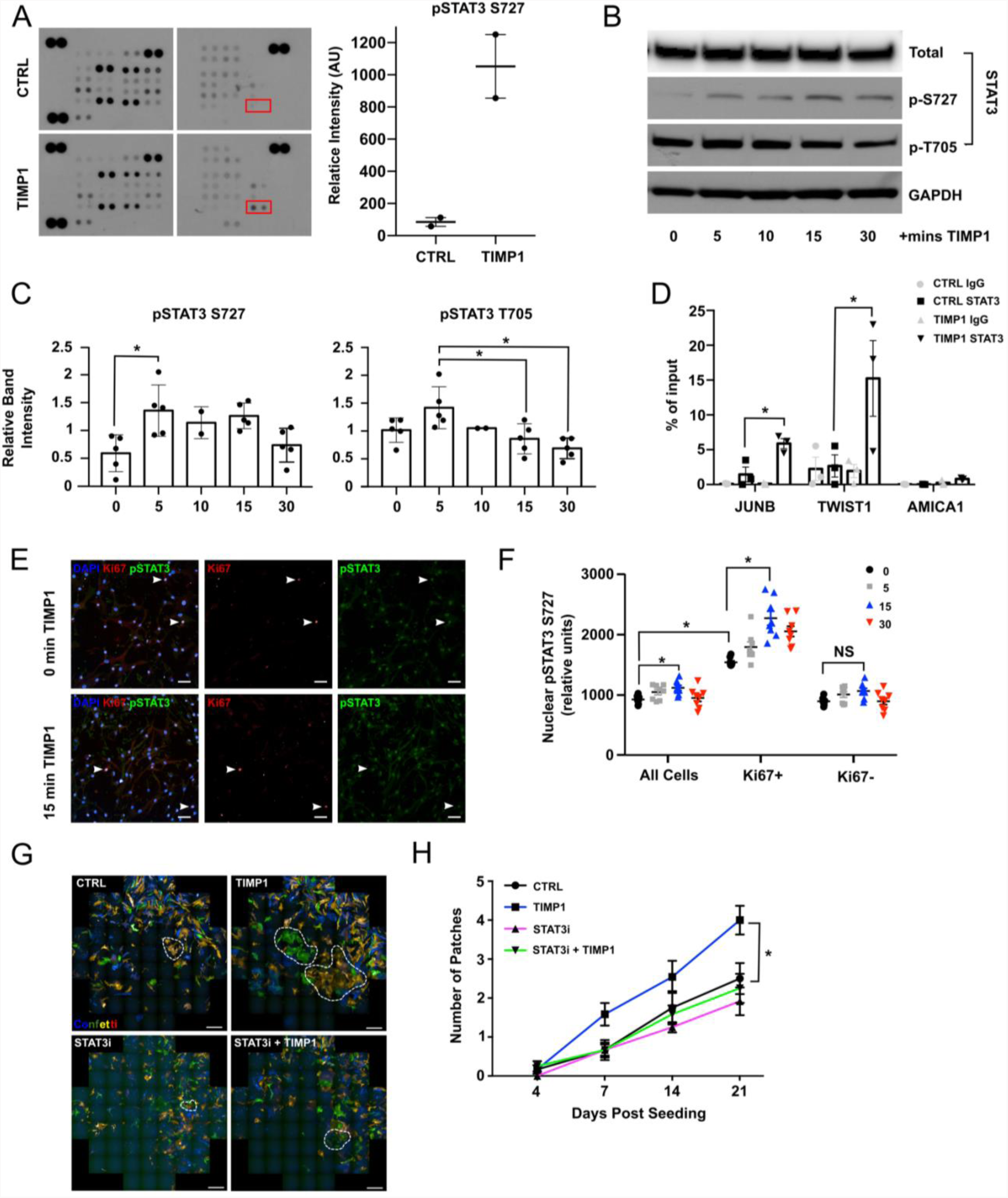
TIMP1 signalling induces STAT3 phosphorylation in human and mouse VSMCs. **(A)** Proteome profiler phosphokinase array of human VSMCs (hVSMCs) following 15 minutes treatment of 500 ng/mL recombinant human (rh) TIMP1 or vehicle control. Quantification of relative spot intensity by densitometry. **(B)** Western blot of total, pS727 and pT705 STAT3 in serum starved hVSMCs analysed 0, 5 10, 15 or 30 minutes after rhTIMP1 treatment. (N=5 hVSMC isolates). **(C)** Quantification of relative band intensity, normalised to total STAT3 levels (N=5 hVSMC isolates), statistical significance determined using ANOVA. **(D)** ChIP qPCR analysis of STAT3 binding at STAT3 targets (*TWIST* and *JUN*) and a negative control (*AMICA1*) gene promoters, in serum starved control hVSMCs and 15 minutes following 500 ng/mL rhTIMP1 treatment. Graph shows anti-STAT3 and control IgG precipitated DNA as a percent of input. N=3 hVSMC isolates. **(E)** Representative images of STAT3 and Ki67 staining in serum starved control hVSMCs and after 15 mins 500 ng/mL rhTIMP1 treatment (N=3 hVSMC isolates analysed in triplicate). Arrowheads indicate KI67+ cells. Scale bar=50 µm. **(F)** Quantification of relative fluorescence intensity of nuclear pSTAT (S727) staining in panel E, for KI67+ and KI67- cells. **(G)** Representative images of lineage labelled VSMCs isolated from Myh11-Confetti aortas cultured 21 days in +/- 500 ng rhTIMP1 and/or 10 uM TT101 (N=6). Scale bar=500 µm. **(H)** Number and size of clonally expanded patches formed by lineage labelled VSMCs from Myh11-Confetti aortas, treated as indicated and imaged over 21 days. Quantified using ImageJ. Statistical significance assessed via generalised linear model. N=6. Asterisk indicates p<0.05.

To assess whether STAT3 phosphorylation is associated with proliferation, we co-stained hVSMCs for KI67 and pSTAT3 (S727) after TIMP1 treatment. Overall, KI67+ cells had higher nuclear STAT3 levels compared to KI67- cells and there was a significantly increased expression 15 minutes after TIMP1 stimulation (**Fig. 6E, F**). There was no effect of TIMP1 on hVSMC proliferation within this time frame (**Extended Data Fig. S7B**). We next analysed VSMCs that were cultured for a short period following isolation from mouse aortas, when acute phenotypic switching and activation of proliferation occurs. STAT3 phosphorylation was also induced by TIMP1 in murine VSMCs cultured for four days, but this effect was less pronounced in cells analysed at day 7 (**Extended Data Fig. S7C**). Interestingly, baseline pSTAT3 intensity was higher in cells cultured for four days after isolation, compared to day 7 cells, suggesting the existence of a “window of opportunity” with higher responsiveness. These data suggest that TIMP1- mediated STAT3 phosphorylation is linked to activation of VSMC proliferation. To directly test this hypothesis, we used a STAT3 inhibitor (TT-101) in the clonal VSMC proliferation assay. TT-101 abolished the increased clone number in response to TIMP1 (**Fig. 6G, H**), demonstrating that STAT3-phosphorylation is important for VSMC proliferation and mediates the effect of TIMP1.

### TIMP1 signals through CD74 to induce VSMC proliferation in a disease relevant mechanism

Signalling by TIMP1 has been reported through several surface receptors that bind to different TIMP1 protein domains (Grünwald *et al*, 2019). We found that the ability to induce clonal VSMC proliferation was retained by the N-terminal part of TIMP1 (**Fig. 7A**), which was recently shown to bind the HLA class II histocompatibility antigen gamma chain, CD74 (Schoeps *et al*, 2021). CD74 is highly expressed in macrophages where it activates pro-survival and proliferation pathways downstream of MIF (Farr *et al*, 2020). In experimental atherosclerosis, CD74 transcription has been detected in modulated, or “macrophage-like” VSMCs (Xue *et al*, 2022). Immunostaining revealed that Myh11-lineage labelled VSMCs also express CD74 protein after vascular injury *in vivo* (**Fig. 7B and Extended Data Fig. S8A**), providing evidence that TIMP1 could function via CD74 in VSMCs. We also detected CD74 in cultured lineage labelled mVSMCs (**Fig. 7C**). We found an association between high pSTAT3 and greater CD74 expression (**Fig. 7C, D**), indicating that CD74 signalling results in phosphorylation of STAT3. To directly test whether TIMP1 acts via CD74 in VSMCs, we used a CD74 blocking antibody, which abolished TIMP1-augmented VSMC clone formation (**Fig. 7E**). Interestingly, anti-CD74 also reduced the frequency of clone formation in untreated cells, which could be due to blocking the effect of TIMP1 secreted by VSMCs after culture-induced phenotypic switching, or other CD74 ligands such as MIF.

**Figure 7.**
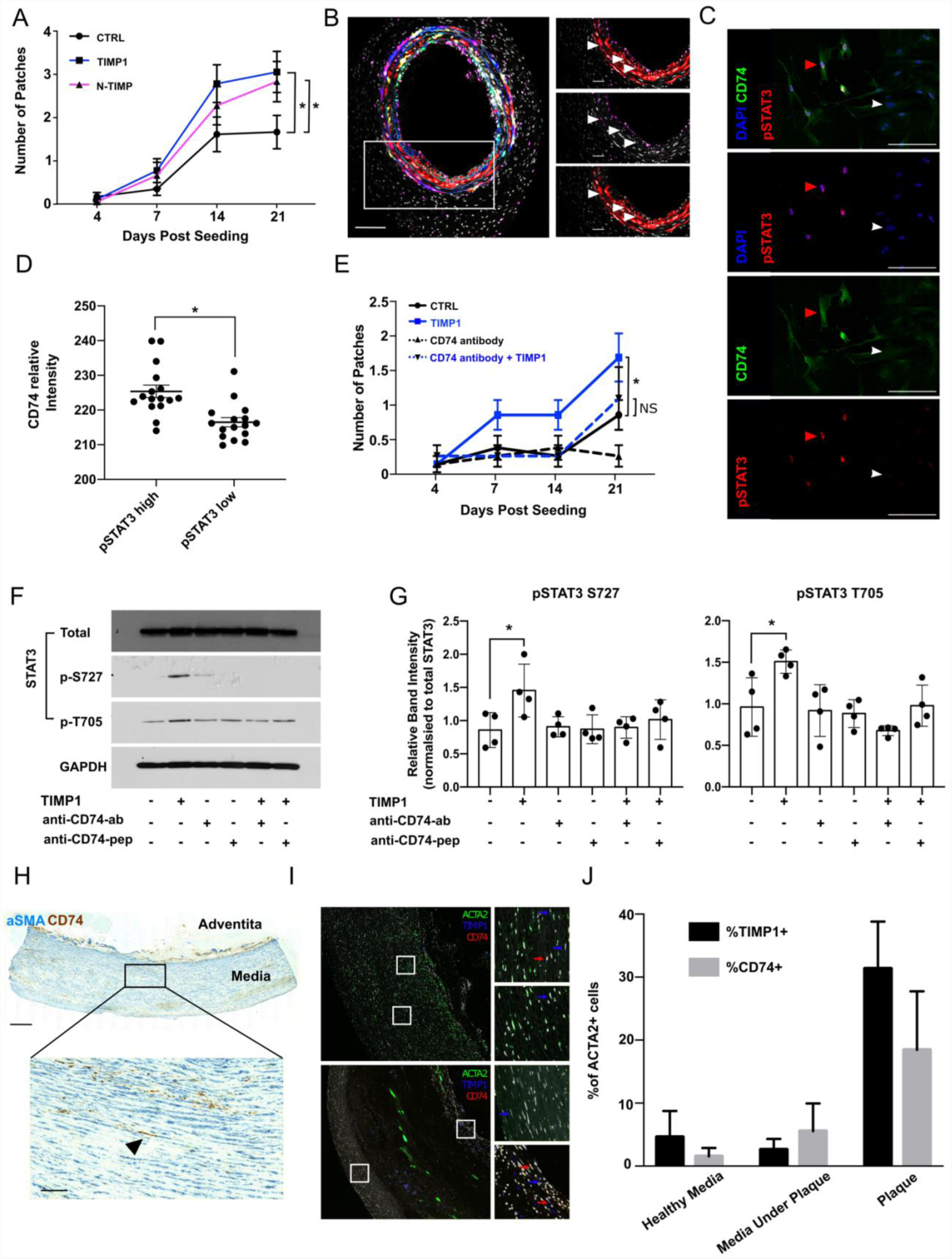
TIMP1 signals to STAT3 via CD74 in a disease relevant mechanism. **(A)** Number and size of clonally expanded patches formed by lineage labelled VSMCs from Myh11-Confetti animals, treated with 500 ng/mL recombinant TIMP1 or equimolar amount of N-TIMP1 tracked over 21 days of culture. Statistical significance assessed via generalised linear model. N=4. **(B)** Representative image of CD74 immunostaining (magenta) in lineage labelled mouse carotids 10 days post ligation of Myh11-Confetti animals. Magnified panels also show DAPI counterstained nuclei (top: merge of Confetti signals, CD74 and DAPI, middle: DAPI and RFP, lower: DAPI and CD74). White pointers marks CD74/RFP double positive cells. N=5 animals. Scale bar=100 µm (overview), 30 µm (zoomed view) **(C)** Representative image of murine VSMCs treated with 500 ng/mL recombinant murine (rm) TIMP1 four days post isolation and immunostained for pSTAT3 (S727, red) and CD74 (green). Nuclei are counterstained with DAPI (blue). Red pointer marks a STAT3 high cells, arrowhead points to a STAT3 low cell in merged and individual channels. **(D)** Quantification of cellular CD74 levels in panel C, stratified by high or low nuclear pSTAT3 intensity. N=4 animals, 4 repeats of each. Statistical significance was determined via student t-test. **(E)** Number of clonally expanded patches formed by lineage labelled VSMCs from Myh11-Confetti animals, treated with 500 ng/mL rmTIMP1 and/or a CD74 blocking antibody or peptide. **(F)** Representative western blot of serum started control human VSMCs (hVSMCs), and cells treated for 5 minutes with 500 ng/mL recombinant human TIMP1 with or without +/- pre-treatment with an antibody (CD74 ab) or a peptide (CD74 pep) that blocks CD74. (**G**) Densitometric quantification of panel F. N=4 hVSMC isolates. (**H**) Representative immunohistochemistry image of CD74 in non-plaque human aorta. αSMA = blue, CD74 = brown. Scale bar=500 µm (overview) and 100 µm (zoomed view). N=7. **(I)** Representative images of RNA *in situ* hybridization for ACTA2 (green), TIMP1 (blue) and CD74 (red) in healthy human aorta, and plaque containing carotid. Blue arrows denote *TIMP1*/*ACTA2*+ cells, red arrows denote *CD74*/*ACTA2*+cells. (**J**) Quantification of the % of *ACTA2* positive cells also expressing *TIMP1* or *CD74* in the medial layer of non-plaque aorta (Healthy media) or in medial (Media Under Plaque) or Plaque regions of carotid endarterectomy samples (N=4 for each condition).

Next, we explored the TIMP1-CD74-pSTAT3 axis in human. As shown in **Fig. 7F**, TIMP1-induced STAT3 phosphorylation was reduced in cells co-treated with the anti-CD74 antibody and a similar effect was observed using a peptide that blocks CD74 action (Figueiredo *et al*, 2018), demonstrating that TIMP1- mediated induction of STAT3-phosphorylation is downstream of CD74 (**Fig. 7 F, G**). We found that CD74 was detected by immunostaining in αSMA+ cells in non-plaque arteries from human organ donors and in samples taken after carotid endartectomies (**Fig. 7H, Extended Data S8B**). To quantify expression, we performed multi-probe *in situ* hybridisation (**Fig. 7I, J**). *TIMP1* and *CD74* transcripts were detected in *ACTA2*+ VSMCs, but observed at low frequency in non-plaque arteries. Elevated frequency of both *CD74*+ (23% of all cells) and *TIMP1*+ cells (16% of all cells) was detected in lesions, and a substantial proportion of these co-expressed *ACTA2* (10-12% of marker positive cells). Jointly, these findings suggest that the induced expression of TIMP1 upon VSMC activation could stimulate VSMC proliferation by binding to CD74 via activating STAT3 phosphorylation.

## Discussion

We here used GRN modelling after acute vascular injury to understand the mechanisms governing activation of VSMC proliferation, which is important to reduce atherosclerosis susceptibility, limit restenosis after vascular intervention and for disease prevention strategies. Our analysis indicates that differential use of shared transcription factors play an important role in this process. This suggests that VSMC state-dependent mechanisms may underlie observed context-specific functions, such as the dichotomous function of the MCP-1 factor, encoded by Ccl2 (Owsiany *et al*, 2022). Using *in silico* simulation experiments, we prioritise factors known to control VSMCs in disease and identify novel candidate regulators with proposed roles in safeguarding the contractile state (e.g. THRA) and promoting cell activation (e.g. RUNX1, CEBP/D, ENO1). We experimentally validate RUNX1 and the TIMP-STAT3-CD74 axis as novel VSMC regulators.

RUNX1 is a pioneer transcription factor that is well-known for its key role in haematopoiesis. Mounting evidence highlights that RUNX1 also impacts differentiation and proliferation in other contexts, including cardiovascular cells (Riddell *et al*, 2020; Koth *et al*, 2020), however, a role in regulating VSMC phenotypic switching has not been examined previously. We here show that RUNX1 increases the frequency of VSMC proliferation, possibly by directly inducing the expression of VSMC “transition state” genes, such as *MMP14*. It is therefore tempting to speculate that this pioneer factor could be assisting the establishment of new transcriptional programs in VSMCs in response to disease risk factors. Our finding that expression of RUNX1 protein in human arteries with intimal thickening is higher compared to those with a healthy morphology suggests that RUNX1 may indeed play a role in vascular changes predisposing to disease.

TIMP1 is a target of the transcriptional rewiring between VSMC states that, unexpectedly, was found to promote establishment of clonal VSMC proliferation. This suggests a positive feedback loop which might impact susceptibility for vascular disease. Interestingly, higher TIMP1 serum levels have been found associated with elevated cardiovascular disease risk in humans (Sundström *et al*, 2004). Future studies are needed to assess whether increased TIMP1 in serum is a consequence of induced TIMP1 expression in activated VSMCs within diseased arteries, or whether elevated circulating TIMP1 functionally increase disease susceptibility by inducing VSMC proliferation and, hence, intimal thickening. In animals, reduced VSMC content and lesion size was observed in *Apoe*-/-, *Timp1*-/- animals compared to Apoe-/- controls, in addition to overall smaller plaque size (Gregoli *et al*, 2016; Silence *et al*, 2002). In contrast, *Timp1*-/- animals had larger neointimal lesions and increased MMP activity following femoral artery injury (Lijnen *et al*, 1999). As VSMC lineage tracing was not performed in these studies, the impact on VSMC proliferation was not directly assessed. Interpreting the effects of global TIMP1 manipulation on VSMC function is further complicated by, firstly, the dual effect of TIMP1 on MMP activity and cell signalling, and, secondly, the ability of TIMP1 signalling to impact multiple cell types in vascular disease. Therefore, targeting of downstream signalling mechanisms would be required to explore therapeutic targeting of TIMP1-mediated VSMC proliferation. In this regard, a recent study showed that TIMP1 affects macrophages via CD74 binding, but that this is dependent on AKT activation (Ebert *et al*, 2023). Here we find that AKT phosphorylation was decreased after TIMP1 treatment in hVSMCs (**Extended Data Fig S6A**). Therefore, it may be possible to exploit the importance of STAT3 phosphorylation for the effect of TIMP1 on VSMC proliferation for specific targeting.

One potential constraint of GRN analyses is the limitation to variably expressed genes. For example, neither the master VSMC regulator myocardin nor its partner SRF show differential expression across the VSMC population at this early timepoint post injury. However, while the motif enrichment analysis in differentially accessible ATAC-seq peaks identified the SRF motif, little change in open chromatin regions at contractile genes was observed, possibly reflecting that epigenetic modification at these SRF targets are not key to the initial VSMC state changes.

We detect an overlap between early VSMC regulators identified here and factors that have been suggested to be important in cultured VSMCs and VSMC-derived plaque cells. For example, enriched motifs in scATAC-seq analysis in human CAD patients include AP-1, RUNX, CEBP, and also highlighted importance of STAT3 in human vascular disease (Turner *et al*, 2022). AP-1 and STAT motifs were also highlighted by trajectory analysis of lineage-traced VSMCs that adopt a more fibroblastic phenotype during mouse atherogenesis (Örd *et al*, 2021). We also detected RUNX1 and CD74 within αSMA+ cells in developed human atherosclerotic lesions. However, whether enrichment of these motifs in atherosclerosis is caused by ongoing VSMC activation in disease, or implies that these factors also play a role at later disease states remains to be tested. Interestingly, studies in zebrafish heart regeneration showed that RUNX1 promotes expression of smooth muscle cell genes in mesenchymal cells (Koth *et al*, 2020), suggesting that in addition to regulating VSMC proliferation, it could also impact fibrous cap formation. The potential mechanistic links between VSMC proliferation and the VSMC-derived cell phenotypes identified in atherosclerotic lesions are an exciting topic for future research.

## Online Methods

### Animals and procedures

All experiments were done according to UK Home Office regulations under project licence P452C9545 and were approved by the Cambridge Animal Welfare and Ethical Review Body. All alleles have been described previously; Myh11-CreERt2 is a Y-linked transgene that confers expression of a tamoxifen-inducible Cre recombinase in smooth muscle cells (Chakraborty *et al*, 2019; Wirth *et al*, 2008), Rosa26-Confetti (Snippert *et al*, 2010) and Rosa26-EYFP (Srinivas *et al*, 2001) are Cre-recombination reporter alleles, KI67-RFP is an insertion in the Mki67 locus resulting in expression of a KI67-RFP fusion protein (Basak *et al*, 2014) and the mutant Apoe allele sensitizes mice to high fat diet (HFD)-induced atherosclerosis development (Piedrahita *et al*, 1992). VSMC lineage labelling was achieved by administration of tamoxifen (10 injections of 1 mg/ml in intraperitoneal injections over two weeks) to Myh11-CreERt2, Rosa26-EYFP (Myh11-EYFP), Myh11- CreERt2, Rosa26-Confetti (Myh11-Confetti), Myh11-CreERt2, Rosa26-Confetti, Apoe-/- (Myh11- Confetti/Apoe), Myh11-CreERt2, Rosa26-EYFP, KI67-RFP (Myh11-EYFP/Mki67-RFP). The tamoxifen administration protocol does not result in equal induction of the four reporter proteins. We find that GFP induction occurs at lower frequency, similar to previous studies (Chappell *et al*, 2016). Myh11-CreERt2 is Y-linked, so all VSMC lineage-tracing experiments were performed using males. Animals were rested for at least 1 week after the last injection to allow tamoxifen washout before tissue harvest, high fat diet (HFD) feeding (21% fat and 0.2% cholesterol, Special Diets Services) or vascular injury. Surgery was performed as described (Chappell *et al*, 2016). The left carotid artery was tied off with a silk suture just under the bifurcation point and analysed 5-28 days after surgery. The animals were given pre-operative analgesic subcutaneously (∼0.1 mg/kg body weight, Buprenorphine) and anaesthetized with isoflurane by inhalation (2.5-3%, 1.5 L/minute for induction, maintained at 1.5%) during surgery. Animals were euthanized by cervical dislocation or CO2 asphyxiation followed by perfusion with cold phosphate buffered saline (PBS) before tissue removal.

### Human tissue

Human arteries were collected after informed consent and ethical approval. Non-plaque aorta was obtained from organ donors via the Cambridge Collaborative Biorepository for Translational Medicine. Carotid endartectomy samples were obtained from the Royal Papworth Hospital research tissue bank. Both male and female tissue and cell isolates were included in all analyses.

### VSMC isolation, culture and treatment

Human VSMCs (hVSMCs) were cultured from aortas of patients undergoing cardiac transplant or aortic valve replacement. After manually removing the endothelial layer and adventitia, the medial layer was cut into 2-3 mm² pieces, placed into 6-well plates containing 1 ml media (DMEM supplemented with 20% fetal calf serum (FCS),100 U/ml penicillin, 100 µg/ml streptomycin) and cultured for 1-2 weeks to allow cells to migrate out of the tissue. After establishment, cells were cultured in hVSMC-specific medium (Promocell, SMC-GM2, C22062) supplemented with 100U/ml penicillin, 100µg/ml streptomycin and were studied at passages 2–10. Human VSMCs were serum starved in media supplemented with 0.1% BSA prior to TIMP1 treatment.

Single cell suspensions of mouse VSMCs (mVSMCs) were generated from freshly isolated aortas of wild type or VSMC lineage-labelled animals (Myh11-Confetti or Myh11-EYFP). Aortas were cleaned off connective tissue, cut open longitudinally and the endothelium removed by gentle scraping with a cotton bud before removal of the adventitia after brief enzymatic digestion. The medial layer was digested to a single-cell suspension in DMEM supplemented with 2.5 mg/mL Collagenase IV (Invitrogen) and 2.5 U/mL Elastase (Worthington) at 37 °C. Cells were cultured in DMEM supplemented with 10% FCS, 100U/ml penicillin, 100µg/ml streptomycin (complete media).

RUNX1 overexpression was achieved by lentiviral transduction. Gibson assembly was used to insert full length mCherry and Runx1 cDNA (*Runx1-202*) linked by T2A into pLentiGFP backbone (Addgene, cat no. 17448). The resulting pLenti-mCherry-Runx1 (or plx302 vector encoding RFP only, empty vector, EV) were co-transfected with third generation lentiviral plasmids (pRRE, pRSV-Rev and pMD2.G) into HEK293FT cells using Trans-iT-LT1 transfection reagent (Mirus MIR2300). Lentivirus-containing medium collected after 48h and 72h after transfection was pooled and concentrated using Lenti-X concentrator (631232, Takara). VSMCs were transduced with lentivirus in media containing 10 µg/ml protamine sulfate (Sigma, P3369). To transiently silence RUNX1, hVSMCs were transfected with 50 nM human *RUNX1*-specific siRNA (ON-TARGETplus® SMART Pool, Dharmacon,L-003926-00-0005) or non-targeting control siRNA (ON-TARGETplus® Control Pool, nontargeting pool, Dharmacon, D-001810-10-05) using Lipofectamine RNAiMAX transfection reagent (13778030, Invitrogen).

Cells were treated with recombinant (r) murine (m) or human (h) TIMP1 (Sino Biological, 50342-MNAH (mouse) and in-house purification as below (murine and human)), N-TIMP1 (murine and human, in-house purification), PDGF-BB (Peprotech, 315-18), GM6001 (Santa Cruz, sc-203979), STAT3i (TTI-101, Cambridge Bioscience, HY-112288), CD74 blocking antibody (clone LN-2, Santa Cruz, sc-6262) or CD74 blocking peptide (Genscript custom, sequence: KSSQSVFYSSNNKNYLA-NH2) as detailed in figure legends. Cells were pre-blocked for 1 hour (CD74 antibody, GM6001, STAT3i) or 6 hours (CD74 peptide) prior to TIMP1 treatment.

### Recombinant TIMP1 protein purification

Recombinant (r) TIMP1 proteins were produced and purified in an endotoxin-free system of HEK293F cells (Thermo Fisher Scientific Inc.) harboring a eukaryotic (pcDNA3.4) expression plasmid encoding the Igkappa antibody chain secretion sequence followed by TIMP1-encoding cDNA sequences (full-length TIMP1 and the N-terminal peptide (N-TIMP1) from both mouse and human, referred to as TIMP1 below), as described previously (Schoeps *et al*, 2021). Cells were cultured in Erlenmeyer flasks at 37°C, 8% CO2, and 125 rpm on an orbital shaker. TIMP1-containing supernatant was purified using an Äktapure Fast Protein Liquid Chromatography (FPLC) system (Cytiva Europe GmbH). The supernatant was applied onto a Capto Q sepharose (Cytiva Europe GmbH) column using the ion exchange chromatography (IEX) equilibration buffer (20 mM HEPES/KOH pH 7.5, 5 mM EDTA, 0.5% Brij35) and the flow through was diluted to a conductivity of maximum 6 mS/cm. Subsequently, it was applied onto a Capto SP sepharose column (Cytiva Europe GmbH) and eluted using the IEX equilibration buffer containing 1 M NaCl. TIMP1- containing fractions were identified by Western blot analysis using the anti-TIMP1 antibody (human: rabbit anti-human antibody; 1:1,000, cat. #8946; Cell Signaling Technology, mouse: goat anti-murine antibody 1:1,000, cat. #AF980; Biotechne R&D biosystems), pooled and concentrated by centrifugal filter (Amicon Ultra 15, 10 kD; Merck). TIMP1-containing concentrates were applied on a HiLoad Superdex 75 pg column (Cytiva Europe GmbH) and eluted by a continuous flow of equilibration buffer (1 × PBS, pH 7.5). Elution fractions containing (N-)TIMP1 were identified by Western blot, and purity was analyzed by Coomassie staining as well as silver staining as described elsewhere. (N-)TIMP1-containing fractions were pooled, concentrated by a centrifugal filter (Amicon Ultra 15, 10 kD; Merck), and stored at −80°C.

### Clonal VSMC proliferation assay

To measure the effect of RUNX1 overexpression on VSMC clonal proliferation, single cell suspensions of medial cells from VSMC-lineage labelled Myh11-EYFP animals were generated, EYFP+ VSMCs isolated by flow cytometry-assisted cell sorting and seeded at a density of 5,000 cells per well of a 96-well imaging plate (CellCarrier-96 Ultra, Perkin Elmer) in DMEM supplemented with 10% (v/v) FBS, 100 U/mL penicillin, 100 mg/mL streptomycin. Low titre lentiviral transduction was performed 3 days post seeding and media was changed twice weekly. For TIMP1 clonal proliferation assays, medial cells from VSMC-lineage labelled Myh11-Confetti animals were mixed with medial cells from wild type animals in a 1:3 ratio and a total of 5,000 cells seeded per well of a 96-well imaging plate (CellCarrier-96 Ultra, Perkin Elmer). Cells were treated as indicated in complete medium from day four after seeding, and medium with fresh reagents was added twice weekly. Cells were imaged 4, 7, 14, and 21 days after plating using an Opera Phenix high content screening system (Perkin Elmer). Image analysis was done using Harmony software (Perkin Elmer) and quantification was performed in Fiji. Patches were defined as an area with three or more contiguous EYFP+RFP+ cells for the RUNX1 experiment and three or more contiguous lineage-labelled cells of the same colour in the TIMP1 assays and patch area was calculated as described (Worssam *et al*, 2022).

### ATAC-seq

Single-cell suspensions of carotid arteries from lineage labelled Myh11-EYFP/Mki67-RFP animals 7 days after carotid ligation surgery and non-ligated littermate controls were generated as described above. Lineage-traced (EYFP+) VSMCs were isolated by cell sorting (BD AriaIII) from non-ligated carotid arteries of control animals. Cells from ligated arteries were stained for SCA1 prior to sorting EYFP+SCA1+ or EYFP+SCA1-cells. The Omni-ATAC protocol was used to process 5,000 cells (Corces *et al*, 2017) from each sample (two independent replicates), using 10-13 PCR amplification cycles and libraries sequenced with a 50 bp paired-end run cycle (Illumina HiSeq or HiSeq2500-RapidRun, 40-60 million reads/sample).

Raw sequence reads were split based on their Nextera barcodes and sequencing quality was assessed using the FASTQC package (Andrews, 2010). Adaptor sequences and base-calls with Phred score < 20 were removed using Trim Galore v.0.6.1, running Cutadapt v.1.18. Reads trimmed to <20bp were discarded and quality-filtered reads aligned to the GRC38.98 Mus musculus genome using Bowtie2 v.2.3.5, discarding reads with >1% error probability. Reads were shifted to account for Tn5 transposase adaptor sequence insertion (+4/-5bp) using alignmentSieve within deeptools v.3.4.3. BAM files were generated using the view command within samtools v.1.18 and deeptools v.3.4.3 was used for conversion to BigWig format for data visualisation in the IGV browser, removing duplicate reads and normalisation by counts per million.

Peak-calling was done after removing reads mapping to a blocklist with repetitive sequences and other mapability issues (a total of 0.02% of the genome, **Supplementary File 3**, samtools, v.1.18) or having low confidence (bedtools v.2.27.1) using MACS2 v.2.2.7.1 (paired-end mode with parameters: –f BAMPE – nomodel –nolambda –q 0.05 –broad). Only peaks that overlapped at least 50% between biological replicates were included in condition-specific peak lists. Peaks were associated with genomic features using ChIPseeker v.1.24.0. A pan-VSMC accessibility list was generated by taking the union of condition-specific peaks.

Each dataset was log2 transformed, normalised (to align peaks with the highest levels of accessibility between samples) and differential accessibility was scored in LIMMA (SeqMonk v.1.47.2) using a >2-fold change threshold and Benjamini and Hochberg adjusted p-value <0.01. For the SCA1+ sample, the top 4000 peaks, ranked by fold-change, were considered in downstream analyses. Peaks were annotated to genes and gene ontology (GO) term enrichment analysis was performed using the online version of GREAT v.4.0.4 (McLean *et al*, 2010) (http://great.stanford.edu/public/html/index.php) with default settings and all genes as a background.

Motif enrichment analysis was done on 500 bp genomic sequences centred on ATAC-seq peak summits (identified using the refinePeak command in MACS2 v.2.2.7.1, Zhang *et al*., 2008), following masking of repetitive elements using RepeatMasker v.4.1.1 (https://www.repeatmasker.org/cgi-bin/WEBRepeatMasker), with default settings using MEME-ChIP v.5.4.1 (Ma *et al*, 2014) for differentially accessible peaks *vs*. a background of all peaks. HOCOMOCO mouse (v.11) was used as the transcription factor motif database, the algorithm was instructed to search for 5 motifs using a CentriMo threshold ≥7. Motifs that were similar to highly repetitive DNA or a very strong true motif, or called as significantly centrally enriched which disagreed with judgement by eye were discarded as spurious.

### Gene regulatory network analysis

Single-cell RNA sequencing (scRNA-seq) profiles of VSMCs isolated from mouse carotid arteries 5 days after carotid ligation surgery (GSE162167) and the bulk ATAC-seq data generated here (accession number below) were used for gene regulatory network (GRN) modelling using CellOracle v.0.10.5 (Kamimoto *et al*, 2023). A “BaseGRN”, including a list of all potential TF-target gene interactions, was constructed using pan-VSMC ATAC-seq peaks. TSS annotation and motif scan steps were performed with this union peak data using the default settings of CellOracle. Cells from the day 5 carotid ligation injury scRNA-seq data, clustered as described in (Worssam *et al*, 2022), were annotated with VSMC states (cluster 9 as CYC, clusters 4 and 3 as PrP, cluster 2 as LNK, clusters 0, 5, 7, 8 as Non-RSP, cluster 1 as stress and clusters 6,10 as path 2 (**Figure 2A, Extended Data Fig. S2A**)) and processed using Scanpy v.1.9.1 (Wolf *et al*, 2018). For GRN modelling, the top 4000 highly variable genes were selected and supplemented with transcription factors that show pseudotime-dependent expression (*Eno1, Id3, Mef2c, Hif1a, Nfia, Hmga1, Ebf1, Rora, Klf9, Jund, Foxp1, Thra, Prrx1, Nfix*), plus *Stat3*, *Twist1*, and *Runx2*.

GRNs were constructed with the top 2,000 interactions for all VSMC states. Topological analyses of GRNs were performed with CellOracle and GRNs were visualized with Cytoscape v.3.7.2 (Shannon *et al*, 2003). GRNs for the stress and path2 states (coloured in grey in Figure 2A) were not investigated further. Venn and Euler diagrams were created by using CRAN R package eulerr v.7.0.0. Gene Ontology (GO) term analysis and visualization were performed with BiNGO v.3.0.4 and GOlorize v.1.0.0.beta1. Communities were detected using clusterMaker2 v.1.3.1 (Morris *et al*, 2011) with the MCODE algorithm (Bader & Hogue, 2003)(fluff option on).

Transcriptional rewiring of nodes was assessed by calculating a rewiring score as the sum of absolute change in connectivity score around each node between the PrP *vs.* Non-RSP networks. Differential gene expression values for cells in PrP *vs.* Non-RSP cell states were calculated using Wilcoxon rank sum-test and gene expression values were plotted using Seurat v.4.3.0 (Hao *et al*, 2021) based on sctransform-normalized values, as described in (Worssam *et al*, 2022). *In silico* simulations were performed with the default settings and top 10,000 interactions in all GRNs with an expression value of zero for the knockout simulations or of 1.5 times the maximum gene expression value for the overexpression simulations.

Prior to GRN modelling with post-carotid ligation day 7 dataset, this data was integrated with the day 5 dataset with sctransform-based normalization using Seurat v.4.3.0 (Hao *et al*, 2021) and further clustered (24 principal components, and 1.9 resolution) in order to identify cell clusters corresponding to equivalent cell states. This information was then used for the GRN modelling.

### Analysis of published human carotid plaque scRNA-seq data

ScRNA-seq profiles (GSE155512) of human atherosclerotic carotid plaques (Pan *et al*, 2020) were filtered to include only cells with the number of genes between 200 and 4,000, total counts more than 500, and percentage of mitochondrial reads less than 10%. The data from three different patients were integrated with sctransform-based normalization and clustered in order to subset VSMCs. Data integration was re-performed with VSMCs only, and the first 17 principal components were used for subsequent analyses with Seurat v.4.3.0 (Hao *et al*, 2021). Differential expression testing was performed for each cell cluster (resolution 0.5) against all other VSMCs with the Wilcoxon Rank Sum test with an adjusted p-value threshold of 0.05. Heatmaps were created with the CRAN R package pheatmap v.1.0.12.

### mRNA isolation and qPCR

RNA was extracted using a RNeasy Mini kit (Qiagen, 74104). cDNA synthesis was performed using QuantiTect Reverse Transcriptase (Qiagen, 205311). Quantitative real-time PCR was performed using SsoAdvanced Universal SYBR Green Supermix (Biorad, 1725270) with the following primer sets. *Mmp14* forward: 5’-GGATGGACACAGAGAACTTCGTG-3’, reverse: 5’-CGAGAGGTAGTTCTGGGTTGAG-3’. *Runx1* forward: 5’-CACCGTCTTTACAAATCCGCCAC-3’, reverse: 5’-CGCTCGGAAAAGGACAAACTCC-3’. *Timp1* forward: 5’-CTCAAGACCTATAGTGCTGGC-3’, reverse: 5’-CAAAGTGACGGCTCTGGTAG-3’.

### Bulk RNA-seq

Bulk RNA-seq was conducted on RNA isolated from of six different hVSMC isolates following 3 hours serum starvation in serum free media containing 0.1% BSA, followed by 6 hours TIMP1 (or vehicle) treatment in serum free media containing 0.1% BSA. Libraries were prepared from oligo-dT-purified mRNA and sequenced on an Illumina Novaseq 6000 (150 bp paired end reads). Raw data reads were trimmed with Trim Galore v.0.6.7, and aligned to the human genome (GRCh38) with Kallisto v.0.46.2. Trimmed Mean of M-values (TMM) normalization was conducted and GSEA performed (Subramanian *et al*, 2005).

### EdU incorporation assay

To assess proliferation, cells seeded in 96-well imaging plates (10,000 per well) were incubated with EdU (10 µM) for 16 hours following treatment as described above (TIMP1/RUNX1 siRNA/OE) and EdU incorporation was detected using the Click-iT EdU kit (C10340, Thermo Fisher Scientific). Briefly, cells were fixed, permeabilized and incubated with the Click-iT reaction cocktail (containing CuSO4, Alexa Fluor 647-conjugated azide, reaction buffer and additive) for 30 minutes protected from light. Next, cells were washed and stained with DAPI for 10 minutes and imaged using an Opera Phenix high content screening system (Perkin Elmer). Quantification of EdU+ cells was done using Harmony software (Perkin Elmer), by thresholding of intensity values measured in detected nuclei.

### Western blotting

Whole cell protein lysates were prepared in RIPA buffer freshly supplemented with proteinase inhibitors (Millipore) and phosphatase inhibitors (Millipore). Protein concentration was determined using the BCA method (23227, Pierce BCA protein assay kit, Thermo Fisher). Immunoblotting was performed according to standard conditions, using gradient (4-12%) polyacrylamide gels, methanol-based wet transfer and chemiluminescence detection (Amersham ECL detection reagent, GE Healthcare). Primary antibodies (total STAT3 (Cell Signaling Technologies, 9139), STAT3 pS727 (Cell Signaling Technologies, 9134), STAT3 pT705 (Cell Signaling Technologies, 9145), and GAPDH (loading control, Cell Signaling Technologies, 2118)) were detected using HRP-labelled secondary antibodies: goat-anti-rabbit (7074S, Cell Signaling Technology).

### Phosphokinase array

Phosphokinase array (R&D Systems, ARY003C) was performed according to manufacturer’s instructions. Densitometry values of spots were calculated in ImageJ and expressed as arbitrary units.

### Immunostaining

Cryosections (14 μm) of ligated left carotid arteries from lineage-labelled Myh11-Confetti animals and plaque containing arteries from Myh11-Confetti/Apoe animals after high fat feeding were permeabilized for 20 minutes in 0.5% (v/v) Triton X-100 (Sigma Aldrich) in PBS. Sections were blocked for 1 hour at room temperature in 1% (w/v) bovine serum albumin and 10% (v/v) of normal goat serum (Dako), and incubated with primary antibody (STAT3 p727 (Abcam, ab86430), CD74 (clone LN1, Biolegend, 151002), KI67 (eBioscience, 14-5698-80)), or isotype control, diluted in blocking buffer overnight at 4°C. Following 3x 5 minutes washes in PBS, Alexa Fluor 647-conjugated secondary antibodies were added for 1 hour at room temperature, sections were washed 3x 5 minutes in PBS and nuclei stained with DAPI (1 μg/mL in PBS, 10 minutes at room temperature) before rinsing in PBS and mounting in RapiClear 1.52 (Sunjin Lab). Confocal imaging was done with a Leica SP8 scanning laser microscope (Leica) using a 20x lens as described (Worssam *et al*, 2022).

Cultured cells were fixed in 4% paraformaldehyde for 10 minutes at RT, washed in PBS, blocked and permeabilised with PBS with 0.3% Triton X-100, 5% goat serum for 1 hour, followed by incubation with primary antibody (STAT3 p727 (Abcam, ab86430), CD74 (clone LN1, Biolegend, 151002), KI67 (eBioscience, 14-5698-80)) diluted in 0.1% Triton PBS 1% BSA for 1 h at room temperature (RT). Cells were washed with PBS, and incubated with the relevant Alexa Fluor 647-conjugated secondary antibody (Invitrogen) diluted in PBS with 0.1% Triton X-100 and 1% BSA for 1 hour at RT. Cells were imaged using an Opera Phenix high content screening system (Perkin Elmer). Image analysis was done using Harmony software (Perkin Elmer), quantification based on intensity values in nuclei defined by DAPI staining, and for mouse VSMCs from Myh11-EYFP animals, within cell borders (EYFP).

Human arteries were formaldehyde-fixed and paraffin-embedded (FFPE) and sections (4 μm) were dewaxed, processed for antigen retrieval and stained as described. Sequential sections were either H&E stained, or co-stained for αSMA (DAKO, M0851), detected with biotin-coupled anti-Mouse (DAKO, E0433) and either RUNX1 (clone A-2, Santa Cruz, sc-365644), KI67 (Cell Signaling Technology, 8114), TIMP1 (clone F31 P2 A5, ThermoFisher, MA1-773), or CD74 (clone LN-2, Santa Cruz, sc-6262) that were detected with HRP-conjugated anti-Rabbit (Cell Signaling Technology, 8114, using DAB peroxidase substrate (SignalStain)).

### Chromatin immunoprecipitation (ChIP)

ChIP was performed on 3×10^6^ hVSMCs using a SimpleChIP kit according to the manufacturer’s instructions (Cell Signaling Technology, 56383). Cells were crosslinked for 15 minutes with formaldehyde (1% final concentration). Chromatin was sheared (15 pulses of 30 seconds with a 30 second rest) using a Diagenode bioruptor and 5 μg of sheared chromatin was immunoprecipitated with 10 μl STAT3 antibody (Clone D3Z2G, Cell Signaling Technology, 12640) or negative control IgG (Cell Signaling Technology, 2729). Protein-G coated magnetic beads were used to capture the antibody-bound protein/DNA complexes. The DNA was eluted, reversed cross-linked, digested with proteinase K and purified.

### FACS and imaging flow cytometry

Single cell suspensions of enzymatically digested medial layer of Myh11-EYFP animals were blocked with TruStain FcX anti-mouse CD16/32 antibody (Biolegend), filtered through a cell strainer (40 μm, Starlab) and sorted (BD FACSAria™ III, BD Bioscience). Positive EYFP signal was determined against an adventitial control sample.

For imaging flow cytometry, single-cell suspensions were incubated with 5 µg/mL TruStain FcX anti-mouse CD16/32 antibody (Biolegend) in FACS buffer (0.5% (w/v) BSA in PBS) for 15 minutes on ice. Cells were then incubated with primary antibody (MMP14 (Abcam, ab53712), SCA1, (Miltenyi, 130-102-343)) for 30 minutes at room temperature and washed twice in FACS buffer. Cells were incubated with secondary antibody in FACS buffer for 30 minutes at room temperature and washed twice in FACS buffer. Intracellular targets were stained using the Foxp3 staining buffer set (eBioscience). Cells were analysed using an Imagestream system (Amnis® ImageStream®XMk II, Luminex).

### RNA scope

RNA *in situ* hybridisation was performed using RNA Scope® Multiplex Fluorescent v2 kits, according to the manufacturer’s instructions (ACD). Experiments used Hs-TIMP1-C2 coupled to opal 570, Hs-CD74-C3 coupled to opal 620 and Hs-ACTA2-C1 coupled to opal 690 (ACD). Imaging was done using a ZEISS Axioscan slide scanner. Analysis was performed in QuPath using the subcellular detection feature.

## Data availability

ScRNA-seq datasets of VSMCs from mouse arteries after injury (Worssam *et al*, 2022) and human carotid plaque cells (Pan *et al*, 2020) are available from the NCBI gene expression omnibus, (GEO, accession numbers GSE162167 and GSE155512). The ATAC-seq datasets and bulk RNA-seq data from TIMP1- treated hVSMCs and controls will be submitted to GEO and accession numbers will be provided. Other data will be made available upon request.

## Acknowledgements

We thank the National Institute for Health Research Cambridge Biomedical Research Centre Cell Phenotyping Hub for cell sorting, Darran Clements at the MRC/Wellcome Stem Cell Institute Advanced Imaging Facility for assistance with confocal imaging, Jo Arnold and Julia Jones at the CRUK for RNA scope hybridisation, the Babraham Institute Sequencing Facility (UKRI-BBSRC Core Capability Grant) and Simon Andrews and Felix Krueger (Babraham Institute’s Bioinformatics) for assistance with ATAC-seq data processing, Jack Palmer and Anirudh Krishnakumar for help with image interpretation and the entire Jørgensen group for advice and discussion. We also thank the donors, their families, the specialist nurses on organ donation (SNOD), and the Papworth Tissue Bank team for human tissue samples. GRN analyses were done using the High Performance Computer cluster at Imperial College Research Computing Service (DOI: 10.14469/hpc/2232). This work was supported by the British Heart Foundation (PG/19/6/34153, FS/15/62/32032, FS/15/38/31516, RM/13/3/30159, RE/13/ 6/30180, RE/18/1/34212, CH/2000003/12800, HFJ and MRB), the Cambridge NIHR Biomedical Research Centre, the Deutsche Forschungsgemeinschaft, Bonn, Germany (KR2047/8-1, KR2047/14-1, and KR2047/15-1 to A.K.), the Medical Research Council of the UK (MC-A652-5QA20, M.S.), and the Chan Zuckerberg Initiative (2018-190766/RG98793 to the Collaborative Bioresource for Translational Medicine).

## Conflicts of Interest Statement

MS is a shareholder of Enhanc3D Genomics Ltd. HFJ is a key opinion leader for Novo Nordisk A/S.

## Notes

### Summary of Updates

Addition of author and correction of errors

