## Supplemental Figures for "Network-based prioritisation and validation of novel regulators of vascular smooth muscle cell proliferation in disease"

**Extended Data**

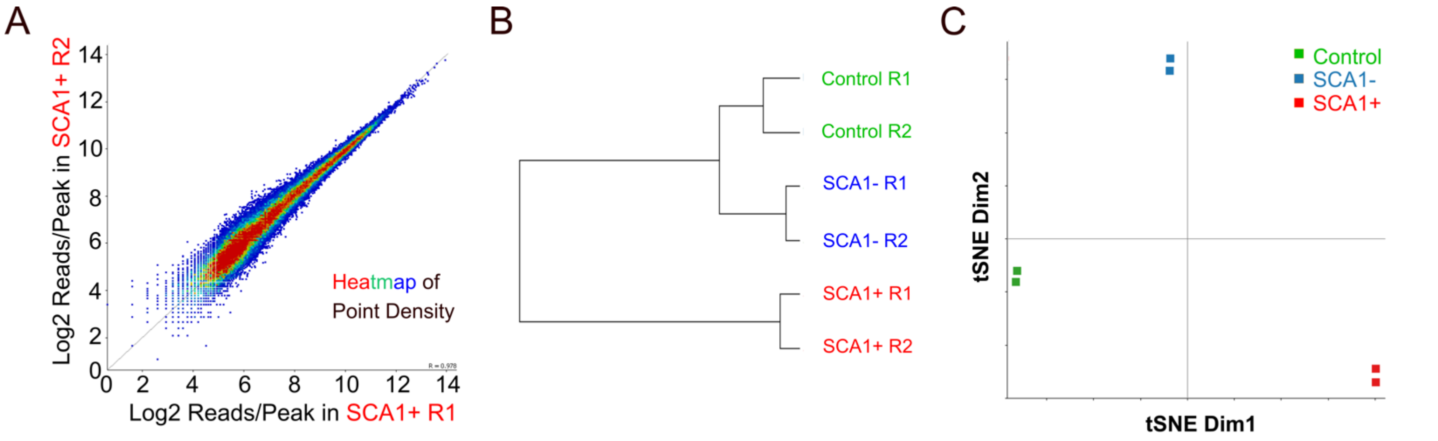


### D GO Biological Process Terms

| Term Name | Binom FDR Q-Val | Binom Fold Enrichment |
|---|---|---|
| muscle cell differentiation | 7.0677e-7 | 2.2033 |
| negative regulation of cell cycle process | 1.2030e-6 | 2.9720 |
| striated muscle cell development | 1.3945e-6 | 3.1915 |
| response to peptide hormone | 2.0448e-6 | 2.3268 |
| striated muscle tissue development | 2.4611e-6 | 2.0468 |
| adrenergic receptor signaling pathway | 3.7598e-6 | 6.2212 |
| regulation of muscle system process | 4.4988e-6 | 2.3743 |
| regulation of actin filament-based process | 7.2077e-6 | 2.1497 |
| cardiac muscle cell differentiation | 7.1800e-6 | 3.1290 |
| positive regulation of organelle organization | 7.1516e-6 | 2.0048 |

### E GO Biological Process Terms

| Term Name | Binom FDR Q-Val | Binom Fold Enrichment |
|---|---|---|
| regulation of cytokine production | 1.6446e-39 | 2.1011 |
| positive regulation of cellular component movement | 2.4561e-37 | 2.0317 |
| positive regulation of cell migration | 6.8387e-37 | 2.0483 |
| positive regulation of cell motility | 2.1819e-36 | 2.0255 |
| inflammatory response | 1.5541e-32 | 2.2837 |
| regulation of cell-cell adhesion | 2.6296e-32 | 2.0920 |
| positive regulation of cell adhesion | 2.2601e-31 | 2.0604 |
| angiogenesis | 7.5796e-31 | 2.0965 |
| positive regulation of cytokine production | 1.3042e-26 | 2.1234 |
| positive regulation of cell-cell adhesion | 7.7041e-25 | 2.2972 |

### F Disease Ontology Terms

| Term Name | Binom FDR Q-Val | Binom Fold Enrichment |
|---|---|---|
| ischemia | 5.1061e-50 | 2.1458 |
| arteriopathy | 8.5213e-47 | 2.2315 |
| myocardial ischemia | 5.3674e-45 | 2.2518 |
| thyroid gland disease | 2.8548e-44 | 2.1500 |
| myeloma | 4.0149e-44 | 2.1129 |
| bone marrow cancer | 9.7460e-44 | 2.0879 |
| bone marrow disease | 1.1249e-43 | 2.0230 |
| hemorrhagic disease | 2.2336e-43 | 2.1441 |
| arteriosclerosis | 2.3423e-43 | 2.2631 |
| arterial occlusive disease | 6.7472e-43 | 2.2421 |

### G MSigDB Signalling Pathway Terms

| Term Name | Binom FDR Q-Val | Binom Fold Enrichmen |
|---|---|---|
| Cytokine-cytokine receptor interaction | 2.1608e-20 | 2.3811 |
| Focal adhesion | 6.4176e-20 | 2.2609 |
| Beta1 integrin cell surface interactions | 1.9436e-16 | 2.9959 |
| Beta3 integrin cell surface interactions | 1.4615e-15 | 3.5247 |
| Natural killer cell mediated cytotoxicity | 2.1745e-15 | 3.0371 |
| Alpha4 beta1 integrin signaling events | 2.8316e-15 | 4.4988 |
| Syndecan-4-mediated signaling events | 4.3179e-15 | 3.9399 |
| ECM-receptor interaction | 1.9599e-14 | 2.5888 |
| Small cell lung cancer | 6.0485e-14 | 2.6296 |
| AP-1 transcription factor network | 1.4853e-13 | 2.7063 |
| Genes involved in Signaling by PDGF | 5.2401e-13 | 2.3354 |
| Renal cell carcinoma | 5.8545e-13 | 2.8130 |
| Jak-STAT signaling pathway | 7.2267e-13 | 2.3082 |
| VEGF signaling pathway | 2.4958e-12 | 3.0655 |

**Figure S1. Quality assessment of ATAC-seq data and gene ontology analysis of differentially accessible genes. (A)** Scatter plot for SCA1+ cells from two independent ATAC-seq experiments showing high replicate correlation. **(B, C)** Hierarchical clustering (B) and principal component analysis (C) of ATAC-seq samples, confirming similarity of replicate samples for all conditions. **(D)** Significantly enriched gene ontology (GO) biological process terms for genes associated with peaks that are more accessible in the control samples from healthy animals, compared to SCA1+ cells isolated from injured arteries. **(E-G)** Significantly enriched GO biological process (E), disease (F) and signalling pathway terms (G) for genes associated with peaks that are more accessible in SCA1+ cells isolated from injured arteries compared to the control samples from healthy animals. Significance was assessed with binomial testing (binom) in the GREAT package.

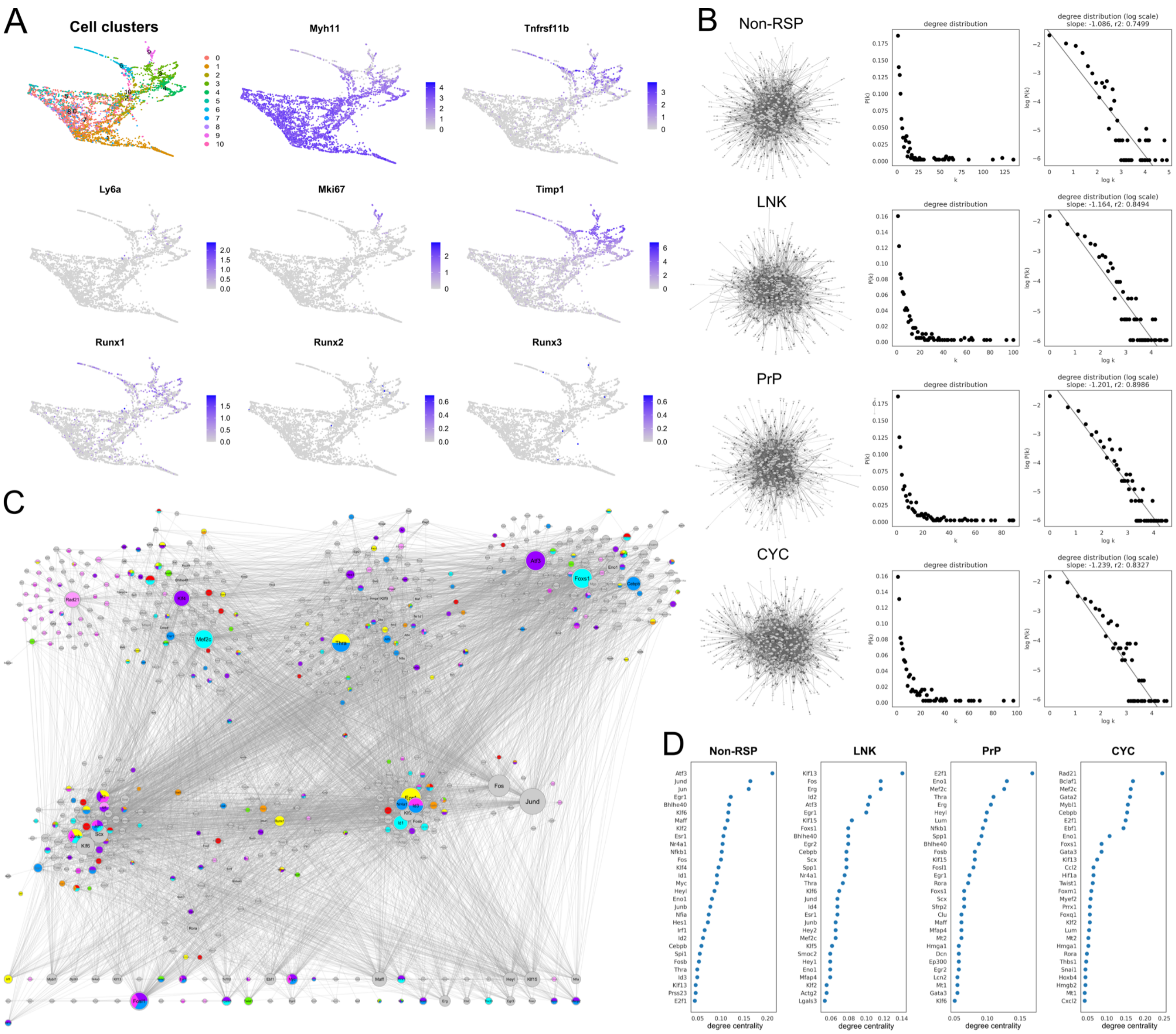


**Figure S2. GRN analysis supplement**

**(A)** Force-directed graph of scRNA-seq profiles of VSMCs analysed 5 days after injury showing the cell clusters and expression of markers characteristic of different VSMC states and selected genes. **(B)** Constructed GRNs and degree distributions. **(C)** Community detection in the union of all four networks, coloured by gene ontology terms as in Figure 2C. **(D)** Nodes with top (30) degree centrality scores for the equivalent GRNs constructed with scRNA-seq profiles of VSMCs analysed 7 days after injury.

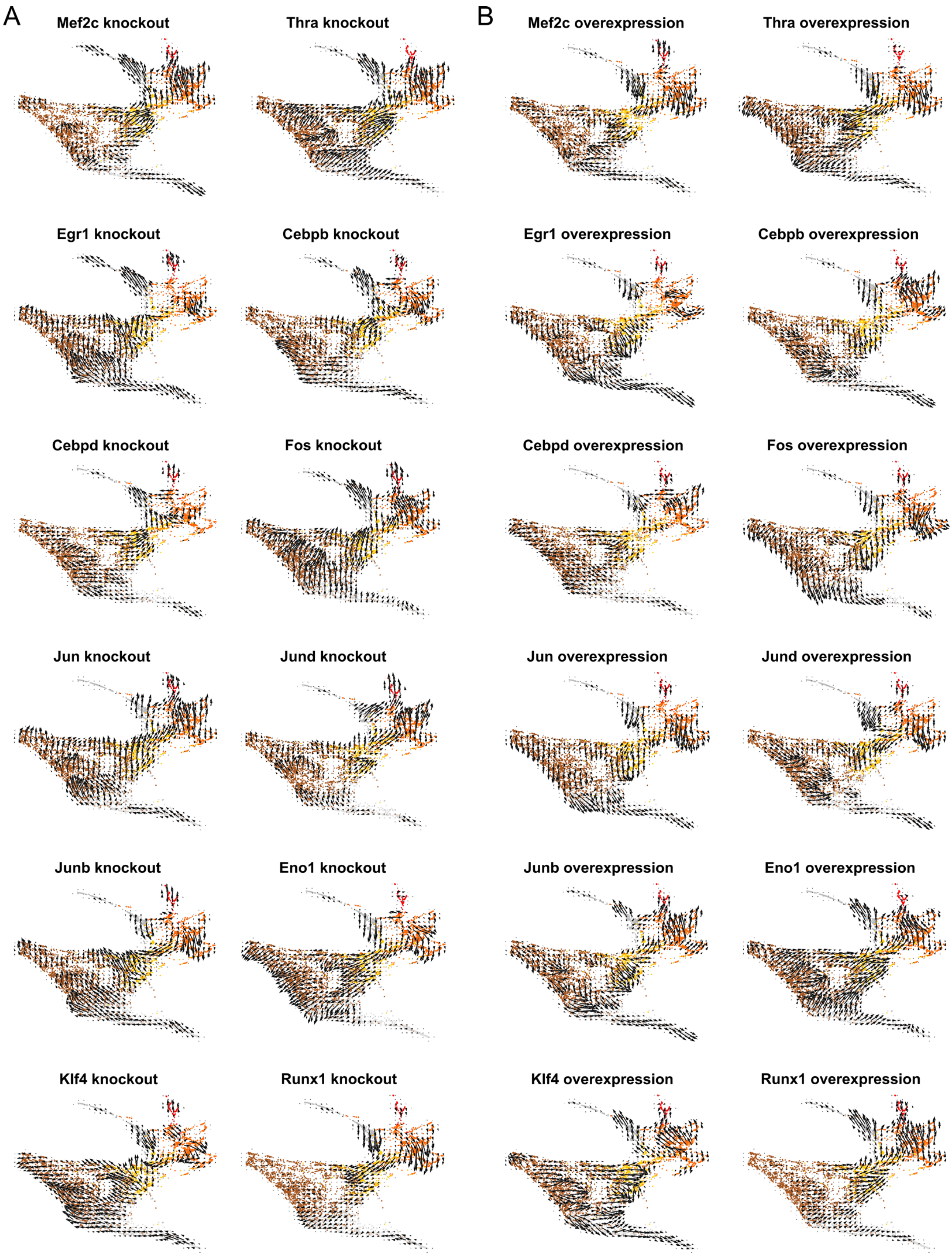


**Figure S3. *In silico* network simulation results**

Force-directed graph of VSMCs isolated 5 days after injury, showing results of *in silico* simulation of knockout **(A)** or overexpression **(B)** simulation for the indicated transcription factors.

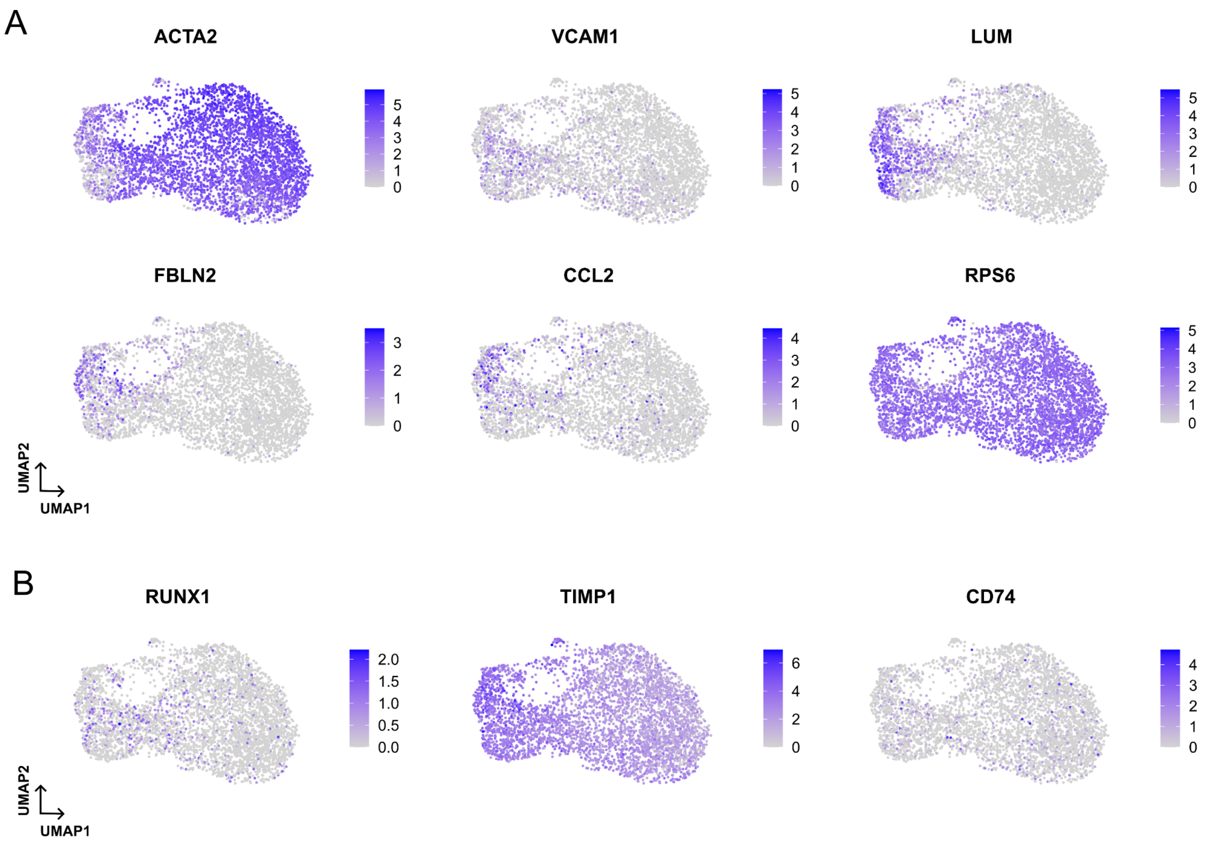


**Figure S4. Gene expression of selected genes in human VSMCs from carotid plaques**

UMAPs showing expression of genes associated with different VSMC clusters **(A)** or of selected genes **(B)** in human carotid plaque scRNA-seq dataset (Pan *et al*. 2020).

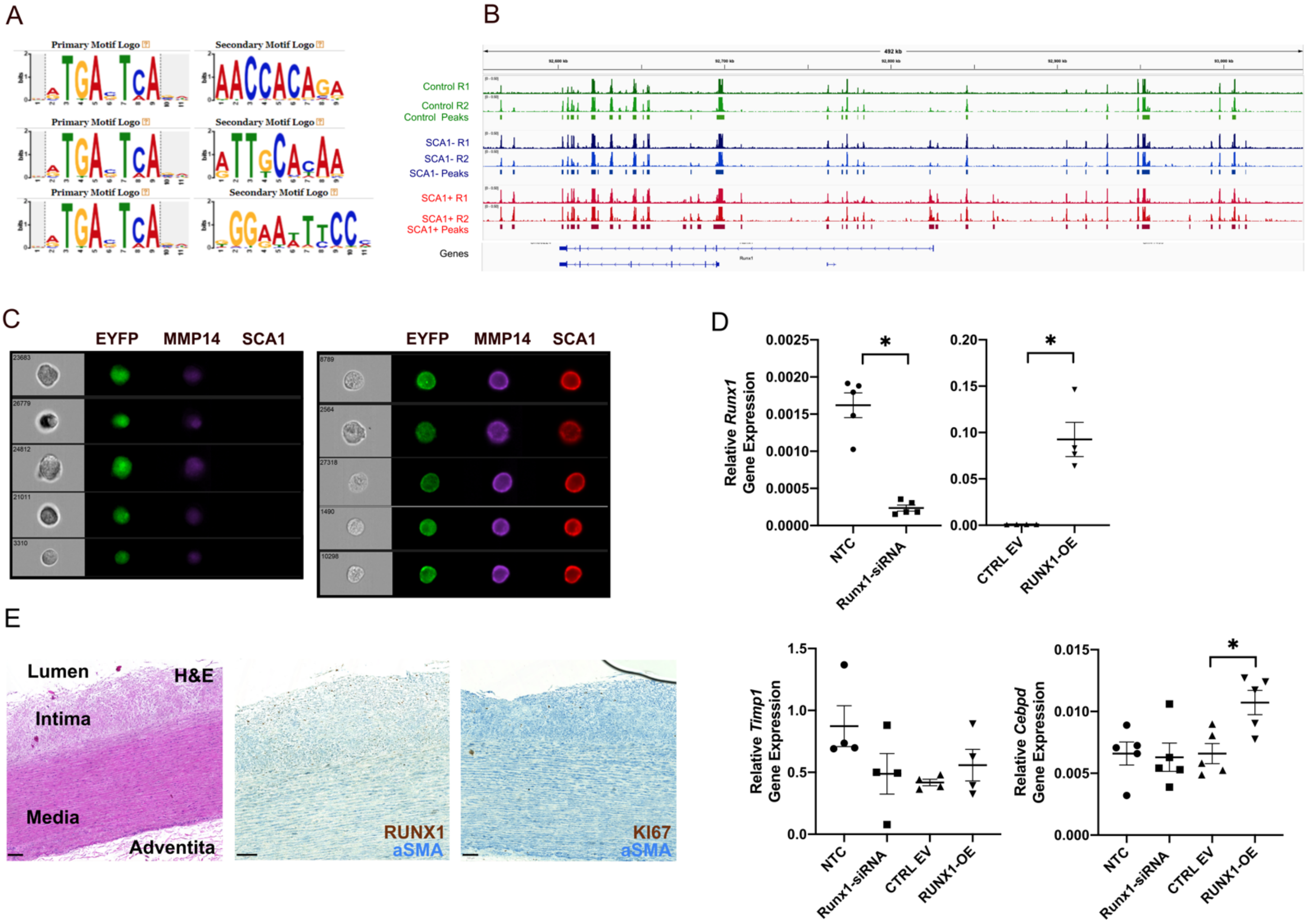


**Figure S5. Analysis of RUNX1 as a candidate VSMC regulator**

**(A)** The RUNX, CEBP and NFκB motifs were detected adjacent to the AP-1 motif in peak showing increased accessibility in SCA1+ cells after injury compared to control VSMCs from no-surgery animals. **(B)** ATAC-seq data tracks for the *Runx1* locus, as described in Figure 1A. **(C)** Imagestream analysis of lineage-labelled EYFP+ (green) VSMCs from Myh11-EYFP animals stained for SCA1 (red) and MMP14 (magenta). Left columns show brightfield images. **(D)** *Runx1, Cebpd* and *Timp1* transcript levels detected by quantitative RT-PCR in mVSMCs transfected with non-targeting (NTC) or *Runx1*-targeting siRNA (Runx1-siRNA), or transduced with an empty vector (CTRL-EV) or RUNX1-overexpressing (OE) lentivirus. Dots represent cells from independent animals (N=5) and lines indicate mean expression. Asterisk indicates $p<0.05$. **(E)** Representative images (N=5) of non-plaque aorta with intimal thickening showing sequential sections after H&E (left) or immunostaining for αSMA (blue) together with either RUNX1 (middle) or KI67 (right) in brown. Scale bars = 100 µm. Lumen, intima, media and adventitia are indicated.

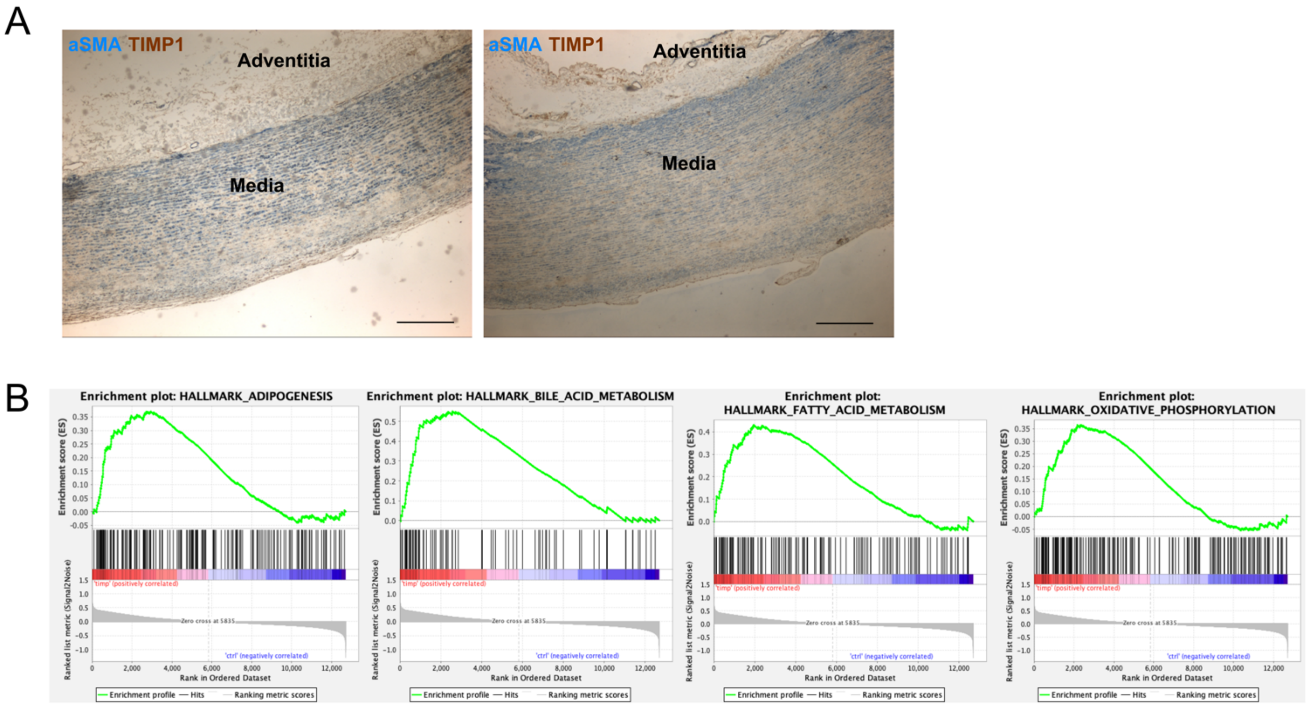


**Figure S6. Impact of TIMP1 on VSMCs**

**(A)** Example immunohistochemistry images for α-smooth muscle actin (αSMA, blue) and TIMP1 (brown) in non-plaque human aorta. Scale bar=500 μm. **(B)** Enrichment plots from gene set enrichment analysis (GSEA) for adipogenesis ($p_{adj}$= 0.023) bile acid metabolism ($p_{adj}$=0.00376), fatty acid metabolism $p_{adj}$=0.00337) and oxidative phosphorylation ($p_{adj}$=0.022) in bulk RNA-seq data from human VSMCs treated with 500 ng/mL recombinant human TIMP1 for 6 hours versus control cells (N=6 independent human VSMC isolates).

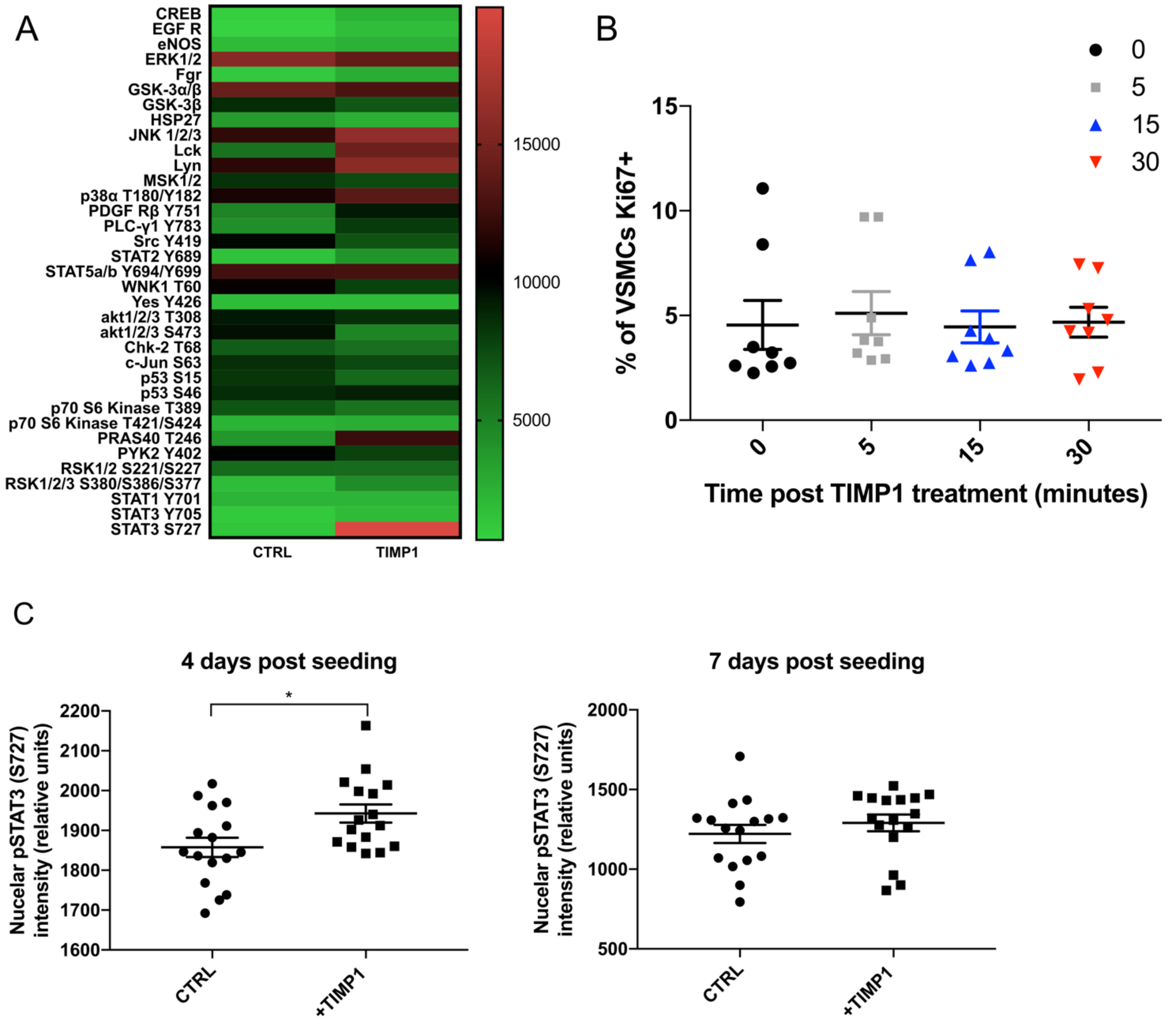


**Figure S7. TIMP1 induces phosphorylation of STAT3 in VSMCs**

**(A)** Heatmap showing quantification of proteome profiler phosphokinase array analysis of human VSMCs (hVSMCs) following 15 minutes treatment with 500 ng/mL recombinant human (rh) TIMP1 or vehicle control. Quantification of relative spot intensity via densitometry. N=1. **(B)** % of KI67+ hVSMCs in samples treated with 500 ng/mL rhTIMP1 following serum starvation and analysed at indicated timepoint. N=3 hVSMC isolates, 3 repeats of each. **(C)** Nuclear intensity of pSTAT3 (S727) in FACS isolated EYFP+ lineage labelled mVSMCs treated with control (CTRL) or 500 ng/mL recombinant murine TIMP1 either 4 (left) or 7 days post seeding (right).

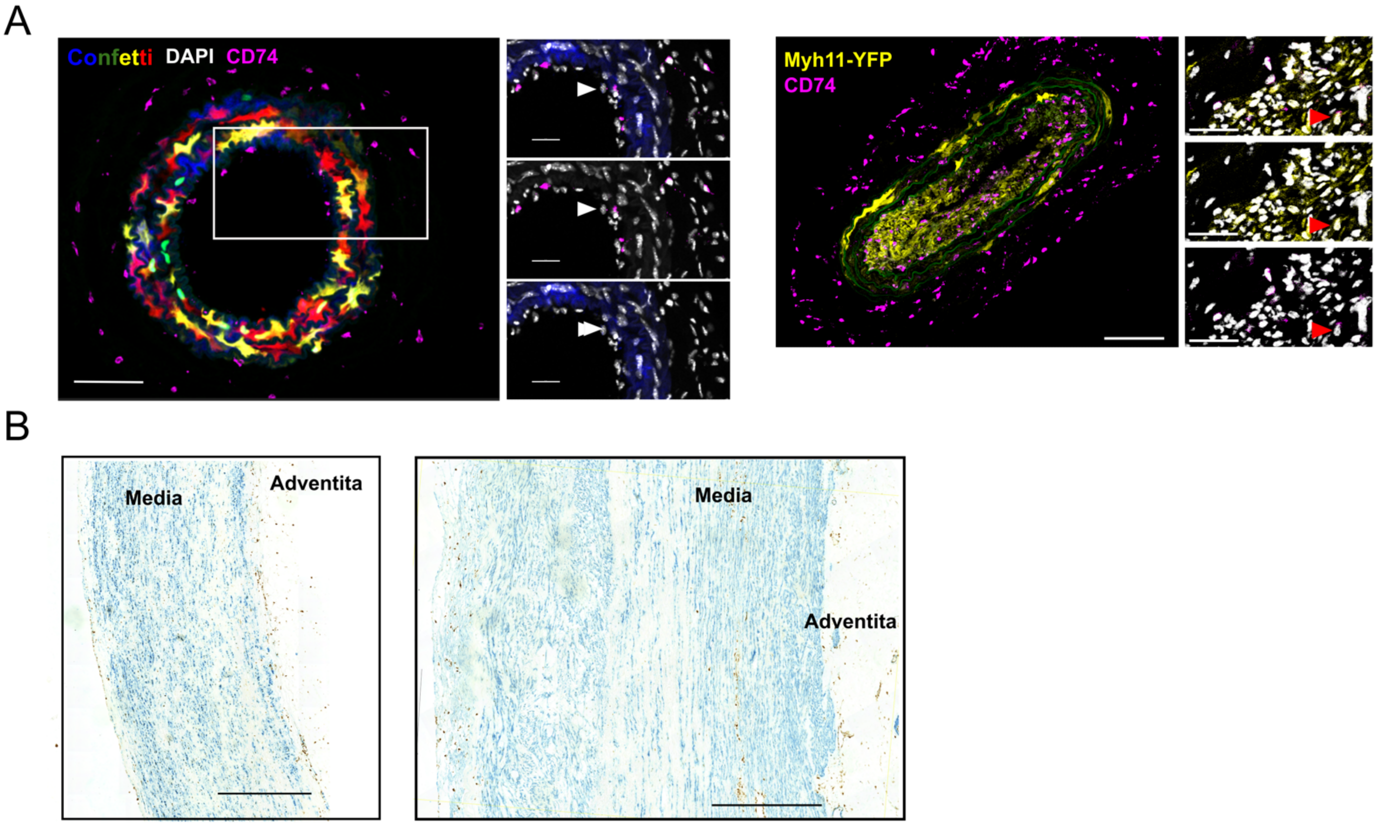


**Figure S8. CD74 expression in VSMCs**
**(A)** Example images of CD74 (magenta) in wholemount images of Myh11-Confetti lineage labelled carotid arteries, analysed 28 days (left) or 10 days post carotid ligation (right). Arrows mark CD74+ lineage labelled VSMCs. Scale bars: 100 µm (overview), 30 µm (zoomed). **(B)** Example images of CD74 (brown) and $\alpha$SMA (blue) immunostaining in non-plaque human aorta. Scale bar=500 µm.
